# Emergent metabolic interactions in resistance to *Clostridioides difficile* invasion

**DOI:** 10.1101/2024.08.29.610284

**Authors:** Achuthan Ambat, Naomi Iris van den Berg, Francisco Zorrilla, Shruti Menon, Abhijit Maji, Arianna Basile, Sudeep Ghimire, Lajos Kalmar, Kiran R. Patil, Joy Scaria

**Affiliations:** Department of Veterinary Pathobiology, Oklahoma State University, USA; Department of Veterinary and Biomedical Sciences, South Dakota State University, USA; MRC Toxicology Unit, University of Cambridge, UK; Department of Bioengineering, Stanford University, USA; Department of Science, Roma Tre University, Italy

## Abstract

Commensal gut bacteria are key contributors to the resilience against pathogen invasion. This is exemplified by the success of fecal microbiota transplantation in treating recurrent *Clostridioides difficile* infection. Yet, characteristics of communities that can confer colonization resistance and the underlying mechanisms remain largely unknown. Here we use a synthetic community of 14 commensal gut bacteria to uncover inter-species interactions and metabolic pathways underpinning the emergent resilience against *C. difficile* invasion. We challenged this synthetic community as well as fecal-matter-derived communities with antibiotic treatment and *C. difficile* in a continuous flow bioreactor. Using generalized Lotka-Volterra and genome-scale metabolic modelling, we identified interactions between *Escherichia coli* and *Bacteroides/Phocaeicola* sp. as key to the pathogen’s suppression. Metabolomics analysis further revealed that fructooligosaccharide metabolism, vitamin B3 biosynthesis, and competition for Stickland metabolism precursors contribute to suppression. Analysis of metagenomics data from patient cohorts and clinical trials attested the *in vivo* relevance of the identified metabolic pathways and the ratio between *Bacteroides* and *Escherichia* in successful colonization resistance. The latter was found to be a much stronger discriminator than commonly used alpha diversity metrics. Our study uncovers emergent microbial interactions in pathogen resistance with implications for rational design of bacteriotherapies.

---

*Clostridioides difficile* infection (CDI) remains a significant challenge to healthcare systems. It has a high incidence in hospital settings due to antibiotic usage which disrupts healthy gut flora and facilitates CDI development and pseudomembranous colitis in humans [1, 2]. *C. difficile’s* spore-forming ability further complicates treatment and contributes to a high recurrence rate, ranging from 13-50%, increasing with each subsequent recurrence [3, 4]. Thus, there is an increasing need for effective prevention and treatment strategies to manage CDI and mitigate its impact on patient health and healthcare systems.

Current preventative measures include pre/probiotics and a fiber-rich diet, which aim to increase gut taxonomic and/or functional diversity. These measures have variable success, and the mechanisms of action, when successful, are not yet fully understood [5]. The clinical practice guidelines recommend oral vancomycin as the primary treatment for non-severe CDI, marking a shift from the earlier preference for metronidazole/vancomycin to vancomycin/fidaxomicin, with fidaxomicin offering the advantage of a lower recurrence rate of about 15–20% [6–13]. The introduction of bezlotoxumab as an adjunctive therapy has been shown to significantly reduce recurrent CDI and maintain a favorable safety profile [14, 15]. The 2013 European Society of Clinical Microbiology and Infectious Diseases (ESCMID-CPG) guidelines recommend metronidazole for non-severe cases and vancomycin for severe CDI [6]. Research into new treatments like ridinilazole as well as the development of vaccines, is ongoing and highlights the advancements in prevention and management of recurrent CDI [16, 17].

Use of antibiotics – of the few that *C. difficile* is not (yet) resistant to – remains a major treatment route but has several adverse side effects, including promoting antibiotic resistance and further disruption of the gut microbial homeostasis [18]. Furthermore, the formation of *C. difficile* spores contributes to antibiotic resistance and the recurrence of infections. Other treatment strategies have been tested such as introducing nutrient competition using nontoxigenic *C. difficile* spores, and modulation of bile acid metabolism for boosting inhibitory compounds like deoxycholic acid and lithocholic acid [19, 20]. The most successful treatment approach yet is Fecal Microbiota Transplantation (FMT) with an overall efficacy rate between 75 and 90% for managing rCDI [21–23]. This treatment is often accompanied by an increase in gut microbial diversity, yet its mechanism(s) of action are not well understood. The success of FMT underscores the emergent resistance provided by the commensal microbiota. Indeed, despite 5% of adults and 15-70% of infants being asymptomatically colonized by *C. difficile*, their healthy microbiome typically suppresses its proliferation/pathogenicity [24]. However, identification of the right donor for a patient remains an unresolved problem due to the lack of mechanistic insights into the resistance conferred by a community and complex interplay between the donor and the host microbiota [25]. Further, FMT carries risks due to the incomplete characterisation of the donor community, including the transmission of infectious agents and immunological side-effects [26]. In severe cases, transmission of multi-drug-resistance organisms have caused US FDA to issue safety alerts [27].

As a safer, more agreeable, and defined treatment option to FMT, bacteriotherapy – administration of a defined microbial cocktail – is emerging as a pre-clinically promising strategy for addressing dysbiosis and combating CDI. However, this strategy has faced challenges in clinical efficacy, as evidenced by trials [28–30]. These outcomes highlight the intricacies of CDI treatment and suggest that a deeper understanding of community ecology, microbial interactions, and associated metabolic outputs is crucial. They also underscore that applying a monotonous treatment strategy to a diverse microbiome signature is often ineffective, calling for more tailored and dynamic therapeutic approaches.

The diversity and dynamics of gut microbiota make it challenging to study the entire microbiome in terms of community ecology and associated colonization resistance mechanisms. To navigate this complexity, we used a synthetic community of 14 bacterial species from a culturomics library [31] that are representative of the core microbiome across populations [32]. Using this defined community, we identify ecological and metabolic interactions underpinning community resistance to *C. difficile* invasion even after antibiotic treatment. Towards assessing the in vivo relevance of our findings, we use fecal-matter derived community as well as perform analysis of all available *C. difficile* cohort metagenomics datasets. Overall, our findings reveal microbial interactions underlying the efficacy of bacteriotherapy and FMT in managing *C. difficile* infection.

## RESULTS

### *Clostridioides difficile* suppression by a 14-member gut bacterial consortium

Fourteen bacterial species – Phocaeicola dorei, Roseburia faecis, Fusicatenibacter saccharivorans, Bacteroides uniformis, Bifidobacterium longum, Blautia obeum, Collinsella aerofaciens, Phocaeicola vulgatus, Parabacteroides merdae, Bacteroides thetaiotaomicron, Bacteroides ovatus, Bacteroides caccae, Escherichia coli, and Parabacteroides distasonis – were selected to assess their impact on C. difficile following antibiotic treatment. These species were previously isolated from a pooled human fecal sample from six healthy donors [31]. The utility of this culture collection is derived from its uniform isolation process, where all bacterial strains were cultured using the same basal medium. Furthermore, these strains were derived from individuals who shared a similar diet and geographic region, ensuring a baseline of prior interaction among them. This uniformity in isolation conditions, coupled with their shared dietary and environmental background, positions these bacterial strains as suitable subjects for studying community ecology within the human gut. To extrapolate the identified community ecology to the diverse gut microbiome population, we selected bacterial species belonging to the core set of microbiome members. The selected 14 microbes represent members of the core human gut microbiota, defined as bacterial species present in 50% of the human gut population with a percentage abundance more than 0.1% [32].

First, we sought to understand the association of these 14 bacterial species with *Clostridioides difficile* infection (CDI) in humans. To this end, we analyzed fecal shotgun metagenomics data from CDI-infected individuals (N=216, across 8 cohort studies), CDI-infected individuals treated with fecal microbiota transplantation (FMT) (N=222, across 6 cohort studies), and healthy individuals (N=251, across 10 cohort studies) from public datasets (Fig. 1a, Suppl. Table 1). In patients with CDI, we observed a significantly lower relative abundance of *P. dorei, R. faecis, B. obeum, F. sacchirovorans, B. uniformis, B. longum, C. aerofaciens, P. vulgatus,* and *P. merdae*. Interestingly, the relative abundance of *E. coli* was significantly higher in CDI patients compared to healthy and FMT-treated individuals (Fig. 1a).

**Fig. 1.**
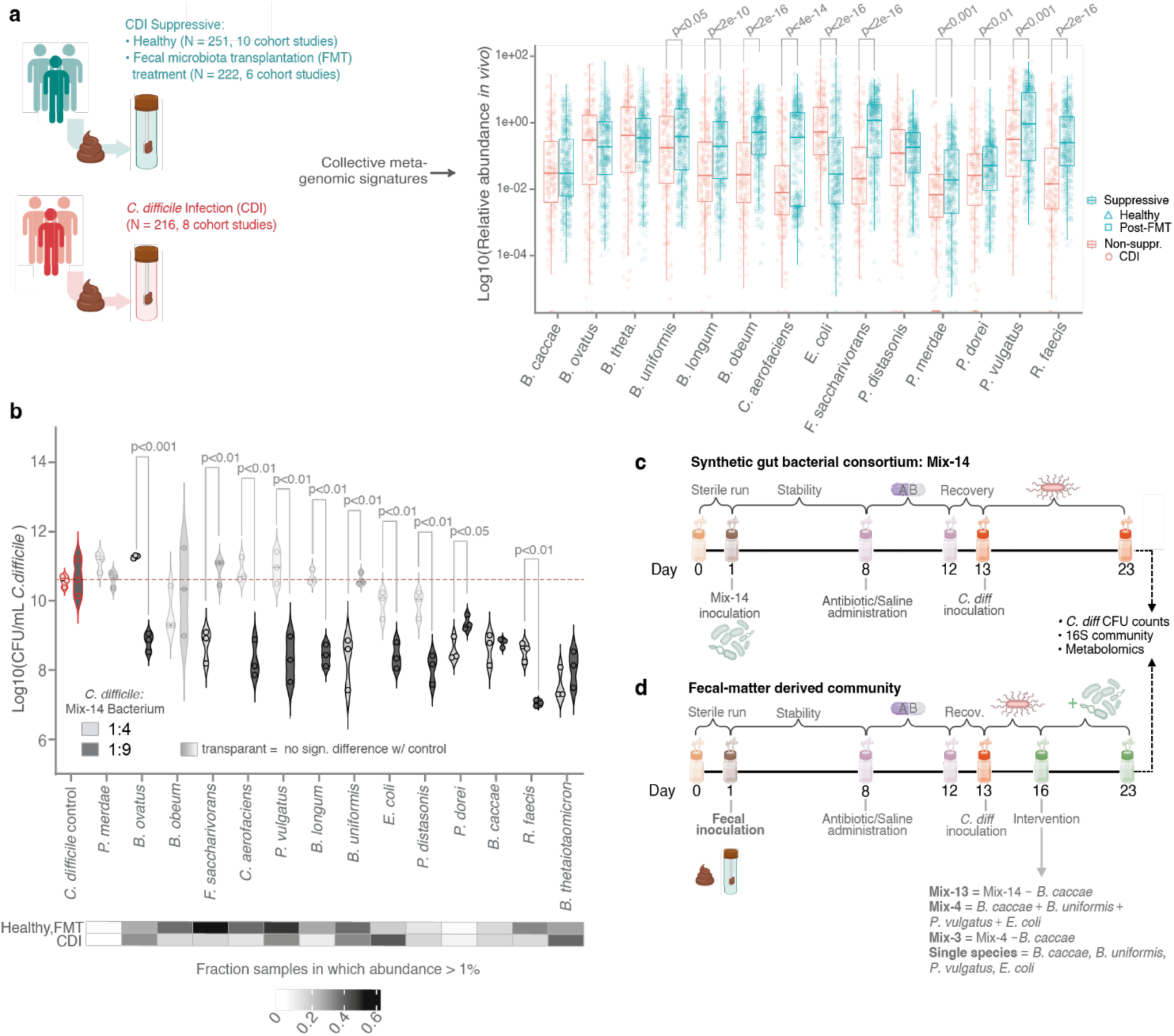
Experimental rationale and design. *C. difficile* suppressive capacities of Mix-14 community members and their differential abundance in CDI patients and healthy subjects. **a.** Relative abundance of Mix-14 consortium species in CDI versus Suppressive (Healthy + FMT-treated) individuals from patient cohort metagenomic data. Most Mix-14 species show a statistically significant differential abundance in healthy versus *C. difficile-*infected patients. Boxplot center lines represent respective medians, box limits represent upper and lower quartiles, and whiskers represent 1.5x interquartile range. **b.** Inhibition of *C. difficile* by 14 different bacterial species in paired co-cultures at two initial inoculum ratios (1:4 and 1:9), representing dysbiotic and non-dysbiotic states, respectively. The heatmap presents each member species’ prevalence across the metagenomic datasets as presented in panel **a**, where prevalence is defined as the fraction of samples where it occupies at least 1% of relative abundance space. **c, d.** Experimental timelines for mapping effects of a synthetic gut bacterial consortium (Mix-14, panel **c**) or fecal-material derived community (panel **d**) on *C. difficile* infection in a continuous bioreactor system. Timeline was designed to mimic typical infection and treatment timeline and duration: e.g., sequential, multiple days of antibiotics; *C. difficile* exposure following antibiotics; and, in the case of fecal-derived communities, bacteriotherapeutic treatment after *C. difficile* infection. Different iterations of the timeline (e.g., with and without antibiotics) were tested. Experimental output included: final *C. difficile* CFU counts, timeseries 16S rDNA amplicon sequencing data, and post-perturbation metabolomics data.

Next, we investigated whether the differential relative abundance of these bacteria between CDI- and non-CDI individuals was related to individual *C. difficile-*inhibitory phenotypes. To address this, we conducted paired co-cultures of each of the 14 member species against *C. difficile.* The co-cultures were conducted under two different initial inoculum ratios; 1:4, representing a higher proportion of pathobiont to gut commensals as observed in dysbiotic states [33–37], and 1:9, corresponding to a lower proportion observed in non-dysbiotic states [38, 39]. Our findings revealed that *P. dorei, B. caccae, R. faecis,* and *B. thetaiotaomicron* were the most effective inhibitors, regardless of the initial inoculum ratio (Fig. 1b). This inhibition was also independent of the individual bacterial generation time, suggesting a critical role of factors other than nutrient competition (Suppl. fig. 2). Most other species exhibited differential inhibition between the initial inoculum ratios tested; with *B. ovatus* even showing a significant boost of *C. difficile* load for the 1:4 inoculum ratio, while showing a significant inhibition for the 1:9 inoculum ratio.

Interestingly, a subset of the strong and consistent individual suppressors, i.e., *B. caccae* and *B. thetaiotaomicron,* did not demonstrate higher prevalence in healthy individuals (Fig. 1a,b). Conversely, species that have a lower relative abundance in CDI patients, e.g., *P. merdae* and *B. obeum,* did not exhibit significant individual suppressive capacity. Collectively, these results indicate that community interactions rather than individual species presence/abundance determine colonization resistance against *C. difficile,* potentially due to taxonomically distinct but functionally equivalent species in healthy and diseased samples carrying out similar metabolic functions.

### *Bacteroides caccae* is a key contributor to *C. difficile* suppression by Mix-14 consortium

Since most of the selected species exhibited a lower differential relative abundance in CDI patient metagenome samples (Fig. 1a), as well as successful individual inhibition of *C. difficile* in at least one of the tested pairwise inoculum ratios (Fig. 1b), we hypothesized that a consortium of these 14 bacterial species could collectively resist *C. difficile* colonization. Given that these 14 bacterial members are prevalent across the healthy population, understanding their dynamics and interactions could help us comprehend the mechanisms behind colonization resistance observed at host-level. Further, to mimic the flow of nutrients in the gut, we used a continuous flow mini bioreactor array (mBRA) model [40] to understand the bacterial interactions between these strains in the presence and absence of abiotic (antibiotic) and biotic (*C. difficile*) perturbations. For this reason, we assembled a consortium of these 14 bacterial members (referred to as Mix-14 hereafter) the bioreactor model system [40]. After establishing a stable community, we administered antibiotics for multiple consecutive days, followed by biotic perturbation via *C. difficile* infection (Fig. 1c). We observed stabilization of community optical density (OD) and pH throughout the experiment for the assembled Mix-14 community in all perturbation regimes studied (Suppl. fig. 1a,b). As hypothesized, Mix-14 successfully inhibited *C. difficile* invasion (i.e., endpoint log10 CFU/mL of < 3.5, i.e., at least five orders of magnitude lower than the *C. difficile* control), with the antibiotic-perturbed Mix-14 showing a more promising inhibition pattern (Fig. 2a).

**Fig. 2.**
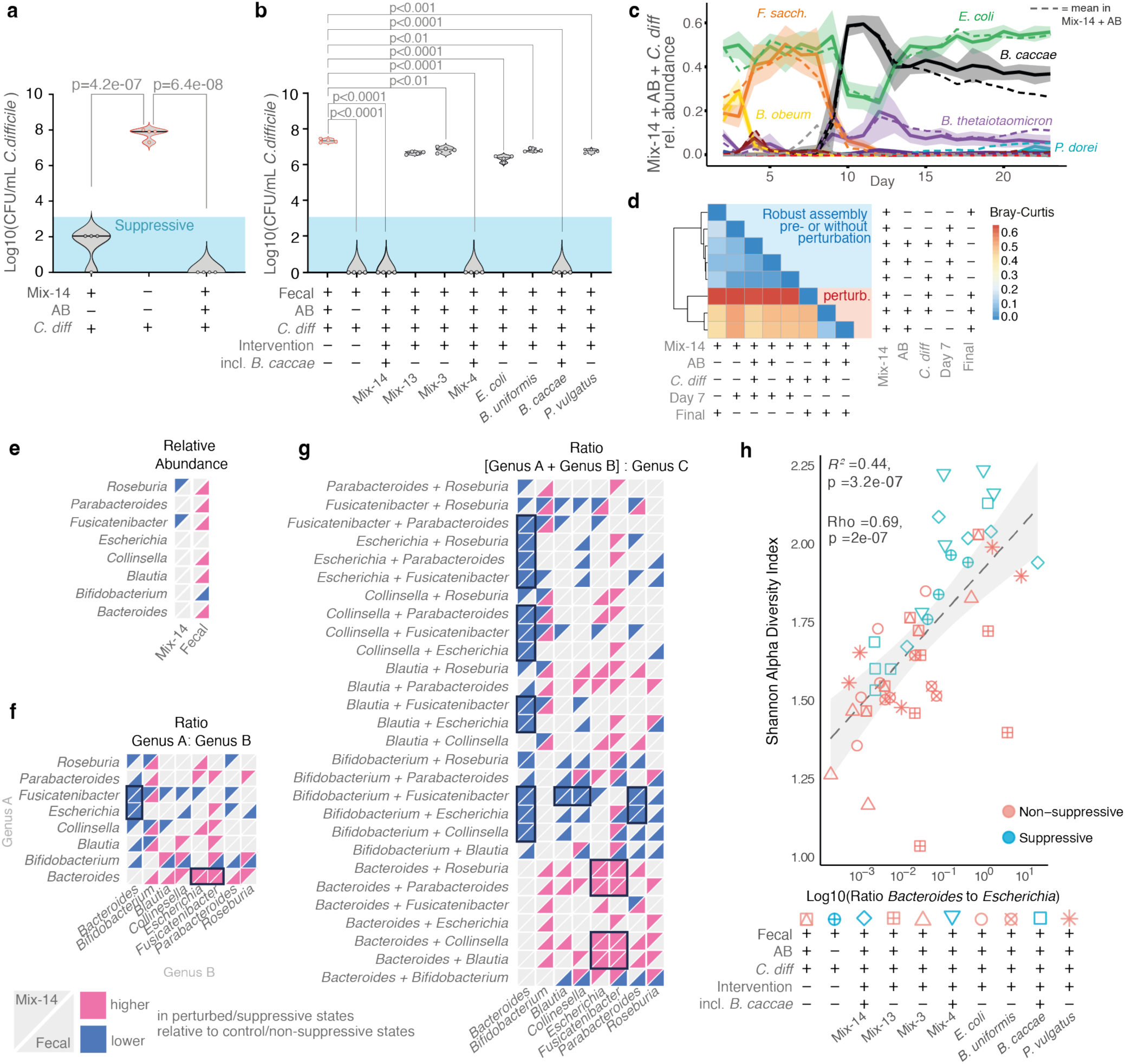
Mix-14 consortium successfully suppresses *C. difficile colonisation in vitro*. **a.** Mix-14 strongly suppressed *C. difficile* with or without preceding antibiotics treatment, lowering final *C. difficile* load by multiple orders of magnitude compared to *C. difficile* control (i.e., *C. difficile* monoculture) at experimental endpoints (day 23). Center lines represent the respective median. **b.** Fecal-matter derived communities suppressed *C. difficile* when not treated with antibiotics, or when treated with antibiotics as well subsequent mixes that contained *B. caccae.* By the experimental endpoint (day 23), suppressive communities successfully lowered *C. difficile* load by several orders of magnitude compared to the non-suppressive fecal community treated with antibiotics (in which final *C. difficile* load was virtually as high as *C. difficile* control as shown in panel **a**). Center lines represent the respective median. **c.** Dynamics of Mix-14 following antibiotics treatment (day 8-11) and *C. difficile* challenge (day 13). While *E. coli* and *F. saccharivorans* were the primary players before antibiotics treatment, both suffer from this perturbation, with *F. saccharivorans* unable to recover. While inhibited by the antibiotics in monoculture, *B. caccae* fills the dynamic niche opened during antibiotic treatment and replaces *F. sacchariavorans* as the key player in an anti-correlative dynamic around system equilibrium with *E. coli*. Following *B. caccae’s* ‘rise’ is *B. thetaiotaomicron*; a key suppressor in monoculture (Fig. 1b). Noteworthy is that the trend of Mix-14 treated with antibiotics but not challenged with *C. difficile* (Mix-14 + AB) was nearly identical (dashed lines), emphasising the robustness of the AB-induced dynamics in Mix-14. Solid lines represent mean trends, and accompanied shaded areas represent ± standard deviation. **d.** Bray-Curtis dissimilarity matrix of Mix-14 pre-perturbation (day 7) and endpoint relative abundance profiles. The matrix shows that all pre-perturbation profiles were similar to each other as well as to the Mix-14 (control) endpoint, emphasising the robustness of community composition and assembly in the absence of perturbations. Post-perturbation profiles were starkly different to the pre- or no perturbation profiles. To illustrate, Mix-14 control’s final community composition is more dissimilar to Mix-14 inoculated with *C. difficile* than Mix-14 inoculated with antibiotics (and *C. difficile*). This suggests not only that the *C. difficile* perturbation induces many (transient) shifts within the community, but also that the perturbations caused by antibiotics mitigate the effects of *C. difficile* inoculation. **e-g.** Relative abundance analyses of Mix-14 and fecal-background communities. Statistical significance (Mann-Whitney U test, BH-adjusted p < 0.05) was determined for genus-level changes in suppressive fecal background communities versus non-suppressive fecal background communities, and perturbed (suppressive) Mix-14 versus non-perturbed (control) Mix-14. Blue indicates statistically significant decline of genus or ratio, while pink indicates a significant increase. For each heatmap cell, the top-half represents trends observed for the defined Mix-14 communities, while the bottom half represents trends observed for the undefined fecal communities. If significance and direction of effect were shared between Mix-14 and fecal communities, respective cells were highlighted with a black outline. In these panels, *Bacteroides* includes *Phocaeicola*, and *Escherichia* includes *Shigella* (Methods). **e.** Mix-14 and fecal-matter derived communities shared no statistically significant genus-level shifts characterising the difference between suppressive and non-suppressive states. **f.** The log10 ratios of *Bacteroides* to *Escherichia* and *Bacteroides* to *Fusicatenibacter* were revealed to be associated with suppressive compositions of Mix-14 and fecal communities. The former ratio had statistical significance of p = 6e-05 in Mix-14, and of p= 0.014 in fecal background communities. The latter had statistical significance of p = 6e-05 in Mix-14, and of p = 0.026 in fecal background communities. **g.** Multiple 2:1 ratios (log10) characterize the difference between suppressive and non-suppressive states in both Mix-14 and fecal communities. Additional key genera were revealed to be associated with potential suppressive internal compositional configurations, namely *Bifidobacterium* and *Collinsella* (e.g., *Bifidobacterium* + *Collinsella* to *Bacteroides,* lower in suppressive communities). **h.** Scatterplot of the logged *Bacteroides* to *Escherichia* ratio (log10, x-axis) versus Shannon alpha diversity (y-axis), calculated for all fecal background communities at day 20. The positive trend between these variables reveals potential related ecological signatures characterising the suppressive communities. The fitted line represents the linear regression, and the shaded area the associated standard error.

To test whether Mix-14 can confer *C. difficile* colonization resistance in a more complex community, we tested Mix-14’s ability to eliminate already-invaded *C. difficile* in a fecal community background. In this setup, instead of initially inoculating with Mix-14, we inoculated the bioreactor wells with the same fecal sample (mix of 6 donors) from which the culture library was created. These communities were then subjected to antibiotics perturbation, followed by *C. difficile* infection and subsequent Mix-14 treatment (Fig. 1d). As hypothesized, Mix-14 intervention successfully eliminated *C. difficile* against a fecal background too (Fig. 2b).

To understand shifts in Mix-14 community composition and dynamics post-perturbation, we analyzed 16S amplicon sequencing data gathered daily over the complete 23-day bioreactor timeline. Our results showed a robust assembly of bacterial communities in all conditions pre- or without perturbation (Fig. 2c, d). Curiously, Mix-14 control’s final community composition was more dissimilar to Mix-14 inoculated with *C. difficile* than Mix-14 inoculated with antibiotics and/or *C. difficile* (Fig. 2d). This suggests not only that *C. difficile* induced many shifts within the community despite being suppressed, but also that the changes in community composition following antibiotics dampen the transient effects of *C. difficile* inoculation.

### Suppressive function and antibiotic response of the consortium is an emergent property

The stabilized Mix-14 community consisted of *E. coli* and *F. saccharivorans* as top-abundant members, followed by *R. faecis, B. thetaiotaomicron, C. aerofaciens, B. obeum,* and *B. caccae,* with the rest of the bacterial species remaining in lower (< 10^-2) or undetectable relative abundance. Interestingly, except for *E. coli*, no other bacterium exhibited monoculture growth that scaled with its final relative abundance in the community. For example, *F. saccharivorans* showed the lowest monoculture growth while being one of the top-abundant species in the community in absence of perturbations, indicating that individual bacterial phenotypes do not necessarily relate to net community performance (Suppl. fig. 2), and further emphasizing the role of community interactions.

Following the antibiotics treatment, we observed a notable shift in community composition and dynamics in Mix-14 compared to the non-perturbed condition (Fig. 2c,d). *E. coli* and *B. caccae* showed the highest relative abundance post-antibiotics. While *E. coli* exhibited a decline in relative abundance during the antibiotics treatment (day 8-11), it recovered once the antibiotics were flushed out of the system, consistent with a bacteriostatic effect of the xenobiotic (Fig. 2c). The sensitivity to antibiotics measured in monoculture did not align with observed community abundance trends. For instance, *B. caccae* strongly increased in relative abundance post antibiotic perturbation, despite showing strong sensitivity to the antibiotics in monoculture (Suppl. fig. 3). This community-level cross-protection is in line with the recent observations in another gut bacterial community [41]. In our study, distinct compositional shifts also emerged after *C. difficile* was introduced without a preceding antibiotics treatment; the Mix-14 community showed an increase in relative abundance for *B. caccae*, along with *B. longum*, *B. obeum*, and *C. aerofaciens,* indicating transient rewiring of community interactions by *C. difficile*.

### Impact of *C. difficile* introduction following antibiotic treatment is synergistic and transient

In the absence of pathogen invasion or antibiotic treatment, the communities assembled robustly, indicating that the observed impact of perturbations is not confounded by variability in the pre-perturbation state of the communities (Fig. 2c,d & Suppl. fig. 4). Since *C. difficile* was suppressed in Mix-14 treated with or without antibiotics, the pathogen’s invasion was transient. *C. difficile’s* short-lived infection of Mix-14 led to significant community shifts, even when not preceded by antibiotics (Fig. 2d). Some species responded similarly to either perturbation, such as *B. caccae* benefitting from both antibiotics as well as *C. difficile* inoculation, while *F. saccharivorans* and *R. faecis* suffered from the abiotic and biotic perturbation.

To further understand the similarities in community-level responses to either or both perturbations across all Mix-14 member species, we constructed a correlation network (Suppl. fig. 6). Within this network, we discerned three distinct clusters. Notably, one cluster comprising *B. thetaiotaomicron*, *P. distasonis*, and *P. dorei* demonstrated a positive covariance post-antibiotic treatment. This cluster, henceforth referred to as the suppressive ecological cluster (SEC), is characterized by its members’ robust individual suppressive abilities (Fig. 1b). The second cluster, made of *F. saccharivorans* and *R. faecis*, positively covaried with each other and negatively covaried with *C. difficile* inoculation and *B. caccae.* In the third cluster, *B. longum, B. obeum* and *C. aerofaciens* positively covaried with each other, yet negatively correlated with *E. coli, B. caccae* and the antibiotics treatment. Yet, this cluster virtually reaches zero relative abundance following antibiotics treatment and was thus considered of negligible importance in driving post-antibiotics *C. difficile* suppression by Mix-14.

### *B. caccae* imparts *C. difficile* resistance to fecal-matter-derived communities

Next, we investigated the degree to which the increase in the relative abundance of *B. caccae* post-antibiotic exposure and/or *C. difficile* infection contributed to colonization resistance. We treated the *C. difficile*-infected fecal community with *B. caccae*, Mix-13 (Mix-14 minus *B. caccae*), and a mock community that is comprised of three species from Mix-14 (i.e., *E. coli*, *B. uniformis*, *P. vulgatus*) based on their diverse monoculture phenotypes (Suppl. fig. 2), with or without *B. caccae* (Mix-4 and 3, respectively). Our results showed that only when *B. caccae* was present in the intervention treatment, the host fecal community exhibited similar colonization resistance compared to the healthy, non-AB treated fecal control (Fig. 2b). Altogether, our results suggest that Mix-14 – both in antibiotics-perturbed and non-antibiotics-perturbed states – could aid community resistance against colonization by *C. difficile*, with *B. caccae* being the key player mediating this inhibition. In accordance with Mix-14 community composition and dynamics, fecal communities displayed extensive shifts in composition as a result of antibiotics treatment (Suppl. fig.7 & 8). Yet, they stabilized following perturbation and were defined by minimal inter-replicate Bray-Curtis dissimilarity between post-perturbation timepoints day 20 and day 23 (Suppl. fig. 7), containing >15 genera with a mean relative abundance of >1% in all conditions (Suppl. fig. 9). These genera included the key genera of Mix-14, with predominant representation of *Bacteroides/Phocaeicola*, Bifidobacterium, and *Escherichia/Shigella* across all conditions. Henceforth, we refer to *Bacteroides* as the taxonomic grouping for both *Bacteroides* and *Phocaeicola*, and refer to *Escherichia* for both *Escherichia* and *Shigella*.

### *Bacteroides* to *Escherichia* ratio underpins suppressiveness of both synthetic and fecal-matter-derived communities

Since perturbed Mix-14 and some fecal communities were *C. difficile*-suppressive, it was of interest to discover which shared taxonomic compositional signatures may be key in conferring suppression. No appreciable genus-level shifts were found to characterize the difference between suppressive and non-suppressive states in both Mix-14 and fecal-background communities (Fig. 2e). Yet, we found that the ratio of *Bacteroides* to *Escherichia* and *Bacteroides* to *Fusicatenibacter* was associated with the composition of suppressive communities shared between the Mix-14 and fecal backgrounds. However, since the latter ratio is an artefact of *Fusicatenibacter* reaching near-zero abundance after perturbation, we did not regard this as a reliable indicator of potential key interactions. The significant discriminatory power of the ratio between *Bacteroides* and *Escherichia* – while either genus alone fails to achieve the same – further supports that the suppressive effect of *Bacteroides caccae* on *C. difficile* is not merely a matter of its abundance but also depends on its proportion within the microbial community, highlighting the complex role of community composition in resistance (Fig. 2f).

We further investigated internal community configurations as potential ‘biomarkers’ of suppressiveness, assessing the complete combinatorial space of bacterial ratios at 2:1 ([Genus A + Genus B] to genus C), allowing the lower-abundant genera to reveal statistical relevance in relation to other genera. Of the 168 ratios thus tested, 24 were found to be significant. In addition to the ratio of *Bacteroides* (+ other genera, such as *Parabacteroides*) to *Escherichia* being significantly higher in perturbed and/or suppressive states of both Mix-14 and fecal background communities, several inverse ratios, e.g., *Escherichia* + *Roseburia* to *Bacteroides* were significantly lower (Fig. 2g). Some of these ratios, such as *Bacteroides* + *Parabacteroides* to *Escherichia,* were significant *despite* the added genus; i.e., the added genus slightly reduces the statistical power of the underlying 1:1 ratio that ‘carries’ the significance. Yet, additional genera outside of this statistical ‘masking’ effect were revealed to be associated with suppressive sub-communities. For instance, in fecal communities, the ratio of *Bifidobacterium* + *Escherichia* to *Bacteroides* was a stronger discriminator (p = 0.0004) than *Escherichia* to *Bacteroides* (p = 0.014). To assess to what extent these ecological signatures of suppression related to higher alpha diversity – the ecological signature typically associated with *C. difficile* suppression [2] – we investigated the correlation between these metrics. The ratio of *Bacteroides* to *Escherichia* correlated significantly with alpha diversity (Fig. 2h, Suppl. Table 2). Altogether, our analysis supports the ratio of *Bacteroides* to *Escherichia* as one of the key ecological signatures of *C. difficile* inhibition *in vitro*, along with higher alpha diversity.

### Genome-scale modelling identifies metabolic interactions connecting *C. difficile* suppressive consortium

Genome scale metabolic models were reconstructed for the Mix-14 consortium species based on the assembled genome sequences and used to assess auxotrophies and potential inter-species dependencies using flux balance analysis (FBA). *F. saccharivorans* featured 11 auxotrophies, including that for riboflavin and benzoate, which might make it more susceptible to environmental variations impacting availability of these nutrients. On the other side, *B. caccae*, which negatively correlated with *F. saccharivorans*, harbours more biosynthetic capabilities, rendering it relatively less dependent on nutritional fluctuations. Notably, *C. difficile* was predicted to be auxotrophic for a range of amino acids, including L-isoleucine and L-tryptophan, indicating susceptibility to competitors for these metabolites. In single-species simulations, both *B. caccae* and *C. difficile* showed a preference for L-isoleucine suggesting competition between the two in the experimental conditions.

Next, we used metabolic models for simulating metabolic competition and cross-feeding within the Mix-14 consortium. The simulations predicted that *E. coli* is a potential donor providing various metabolites to eight community members, as well as *C. difficile* (Fig. 3a). Consistent with its auxotrophies, *C. difficile* was predicted to be dependent on the supply of isoleucine and tryptophan from other community members (Methods). Both amino acids are Stickland fermentation substrates supporting the growth for *C. difficile* [1]. This metabolic role of *E. coli’s* suggests that its contribution to Mix-14 in colonization resistance goes beyond competitive exclusions despite its dominance in the community.

**Fig. 3.**
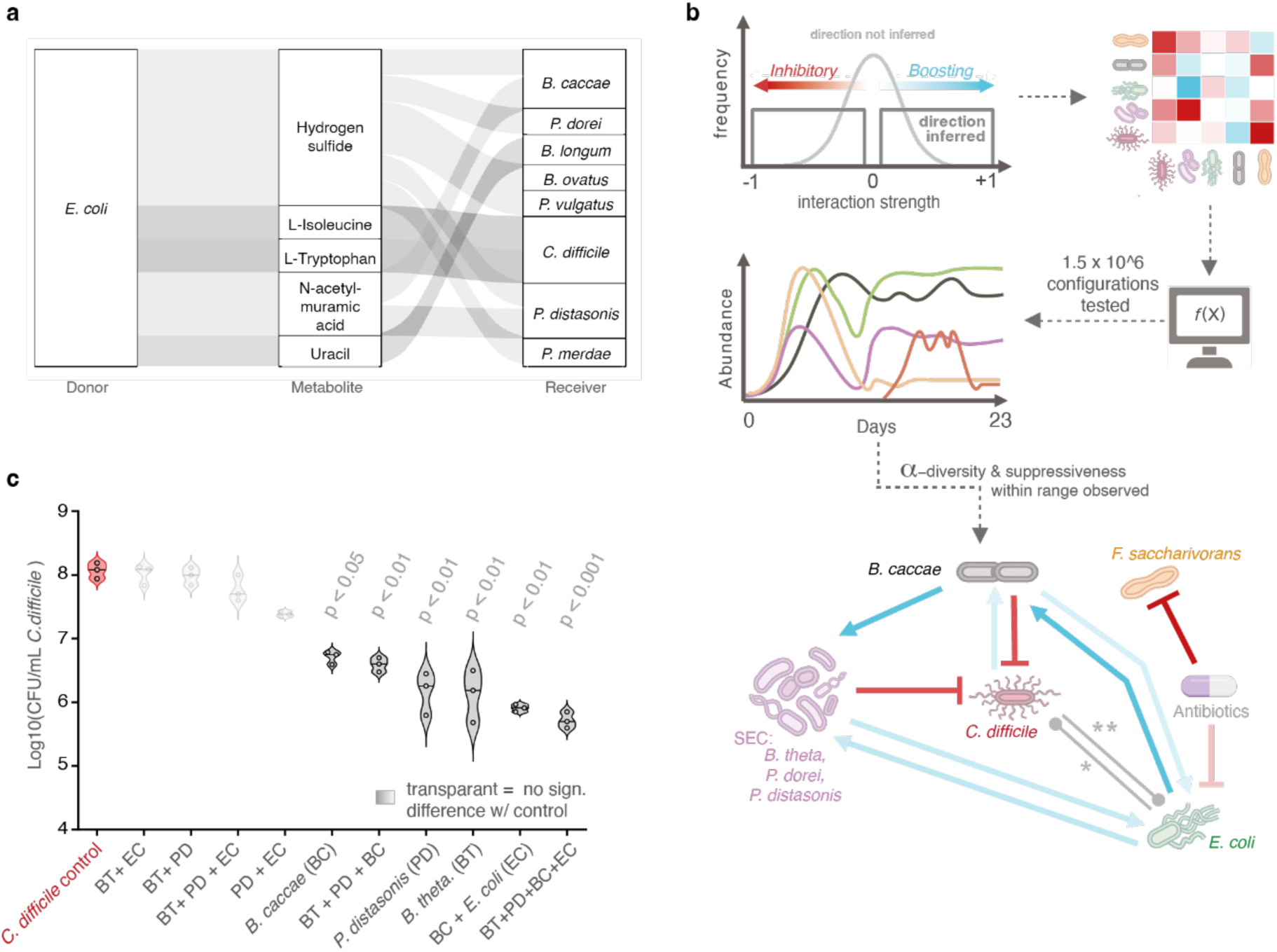
Genome-scale and ecological models indicate key interactions amongst Mix-14 community members. **a.** Alluvial diagram highlighting the predicted nutritional dependencies in Mix-14 based on metabolic models and flux balance analysis-based methods. Of note is the central donor role of *E. coli* to several key Mix-14 members, as well as *C. difficile*. The thickness of each flow is proportional to the average SMETANA score across simulations, and only interactions with a score greater than 0.75 are shown. **b.** Mix-14 *in silico* experimentation pipeline, resulting in a proposed Mix-14 interaction network. Key interactions (i.e., those most likely to reproduce observed community dynamics and functions) were deduced from over a million simulation results. In the shown network containing key interactions, arrow opacity corresponds with predicted interaction strength (the fainter, the weaker), while arrow colour corresponds with predicted net interaction direction (red = inhibitory, blue = boosting). * Model results predict that this interaction type could range from weakly positive to moderately negative without compromising reproducibility of Mix-14’s observed suppressiveness and community composition. While *E. coli* was observed to inhibit *C. difficile* in a paired co-culture (for inoculation ratio of 1:9, Fig. 1b), its net effect within the community co-culture may be different (e.g., *E. coli* secreting different compounds in the medium, see GEM-predicted nutritional dependencies in panel **a**). ** Model results predict that Mix-14’s suppressiveness could be maintained if *C. difficile* inhibited *E. coli* weakly or moderately. If strongly, suppressiveness as well as accuracy in reproducing Shannon alpha diversity was lost. A neutral or positive effect of *C. difficile* on *E. coli* sustained predicted suppressiveness. **c.** Experiment validation of model predictions on the role of sub-consortia in Mix-14 in conferring C. difficile suppressiveness. For example, only when paired with B. caccae did the combination of SEC-species B. thetaiotaomicron and P. distasonis show significant suppressive capacity (final C. difficile CFU/mL >2 orders of magnitude lower than control). The suppressive effect was most pronounced when B. caccae and E. coli co-occurred, with the duo nearly mirroring this suppressive capacity independently. This finding underscores the critical role of E. coli in bolstering B. caccae’s net suppressive ability, highlighting the significance of their interaction within the Mix-14 community. Center lines represent respective medians.

### Ecological modelling uncovers interaction between *Bacteroides* and *Escherichia* driving *C. difficile* suppression

Our data shows *C. difficile* suppression to be an emergent, community-scale property rather than the result of a direct/pairwise and isolated interaction between *C. difficile* and any resident community member. To uncover the ecological network underpinning this emergent phenomenon, we used an ODE-based modelling approach. We first inferred and/or constrained the net interactions between community members using available *in vitro* experimental data, including the carbon utilization patterns (95 C-sources) at species and community-scale (Biolog, Suppl. fig. 10), correlation networks within the relative abundance data (Suppl. fig. 6), community final ODs, individual inhibition phenotypes (Fig. 1b), as well as *in silico* data generated via genome-scale metabolic modelling (Fig. 3a). For instance, the anti-correlative behavior around system equilibrium between the main community members *E. coli* and *F. saccharivorans* (without antibiotics) or *B. caccae* (with antibiotics) as seen in Fig. 2c suggests some type of stable co-existence or non-antagonistic interaction. See Suppl. Table 3B for complete details of interaction inference and constraints used in the model.

However, not all sources of data used for interaction type inference agree on (or provide information for) each respective interaction direction or strength. Hence, to explore the variability thereof, we employed a generalized Lotka-Volterra-based model to test how different configurations of the interaction landscape, explored via random permutation (i.e., 1.5 million iterations tested), could reproduce composition distributions, Shannon alpha diversity and *C. difficile* suppressiveness observed for Mix-14 *in vitro* (Methods 4.2.2., Fig. 3b, Suppl. fig. 4 & 5).

The modelling results reveal that, to achieve an *in silico* community capable of reproducing observed suppressiveness, an interaction landscape with a positive effect of *E. coli* on *B. caccae,* and in turn of *B. caccae* on the suppressing SEC-species, was key (Suppl. fig. 11a). In order to achieve observed alpha diversity, the model predicts the interaction landscape is also defined by a positive effect of *E. coli* on the SEC-species (Suppl. fig. 11b), which would correspond with the strong correlations between the end-point relative abundances of these groups (Suppl. fig. 6). Fig. 3b shows key interactions within Mix-14 that the model results predict to represent observed community ecology and associated community outcomes. The importance of the predicted key positive pairing between *E. coli, B. caccae,* and SEC-species corresponds with findings observed for fecal communities (Fig. 2 e-h), and were subsequently validated *in vitro* (Fig. 3c). For instance, only when paired with *B. caccae* did the combination of SEC-species *B. thetaiotaomicron* and *P. distasonis* show significant suppressive capacity (final *C. difficile* CFU/mL >2 orders of magnitude lower than control, Fig. 3c). This suppressive capacity was strongest when *B. caccae* and *E. coli* were also present, yet their paired combination alone could closely approximate this suppressive capacity (also final *C. difficile* CFU/mL >2 orders of magnitude lower than control, Fig. 3c), emphasizing the enhancement of *B. caccae’s* suppressive potential by *E. coli*. Altogether, the ecological modelling results indicate the importance of the interaction between *B. caccae* and the SEC-species, as well as the interaction between *E. coli* and *B. caccae,* in conferring community suppressiveness.

### Metabolomics reveals competitive and inhibitory molecules associated with *C. difficile* suppression

To hunt for molecular mechanisms associated with *Bacteroides*-mediated suppressiveness shared between Mix-14 and fecal communities, we used comprehensive metabolomics analysis and thereby contrasted molecular phenotypes between metabolic landscapes considered suppressive and non-suppressive. Mix-14 + AB, Mix-14 + AB + *C. difficile*, Fecal, Fecal + AB + *C. difficile* + *B. caccae*, Fecal + AB + *C. difficile* + Mix-14 were conditions considered suppressive, while control Mix-14 community, Fecal + AB, Fecal + AB + *C. difficile* were considered non-suppressive. Combining data derived from defined and undefined communities adds considerable background ‘noise’ and other sources of variability that may impede useful statistical comparisons. Hence, to increase replicate space and enhance the ability to find statistically significant trends in the metabolomics data, we used the previously found compositional signatures associated with suppressiveness to guide the extrapolation of suppressiveness capacities of metabolic landscapes belonging to experimental conditions unchallenged by *C. difficile.* For instance, we considered the supernatant of fecal control as metabolically ‘suppressive’, since the condition ‘fecal + *C. difficile’* was suppressive. Similarly, we considered the supernatant of ‘Mix-14 + antibiotics’ as ‘suppressive’, since ‘Mix-14 + antibiotics + *C. difficile*’ was suppressive. This method of extrapolation may introduce false positive or negative assignments of metabolites as suppression-associated, however, since community compositions of extrapolated communities were very similar to their *C. difficile*–challenged counterparts, this limitation was considered of negligible importance.

We found a total of 47 metabolites with significant differential abundance (Wilcoxon rank sum test, BH adjusted p-value < 0.05) between *C. difficile*-suppressive and non-suppressive samples (Fig. 4a). Out of these 47 metabolites, 36 were depleted in suppressive samples, and 11 were enriched. Out of the 36 depleted metabolites, 16 corresponded to amino acids and their derivatives, hinting at the importance of these substrates for *C. difficile*’s growth via Stickland metabolism. This is in line with previous metabolomics studies on *C. difficile* pathogenesis. For example, enrichment of lipids and metabolic byproducts during *C. difficile* colonization was previously noted [42], and accordingly, these metabolites were depleted in our suppressive samples.

**Fig. 4.**
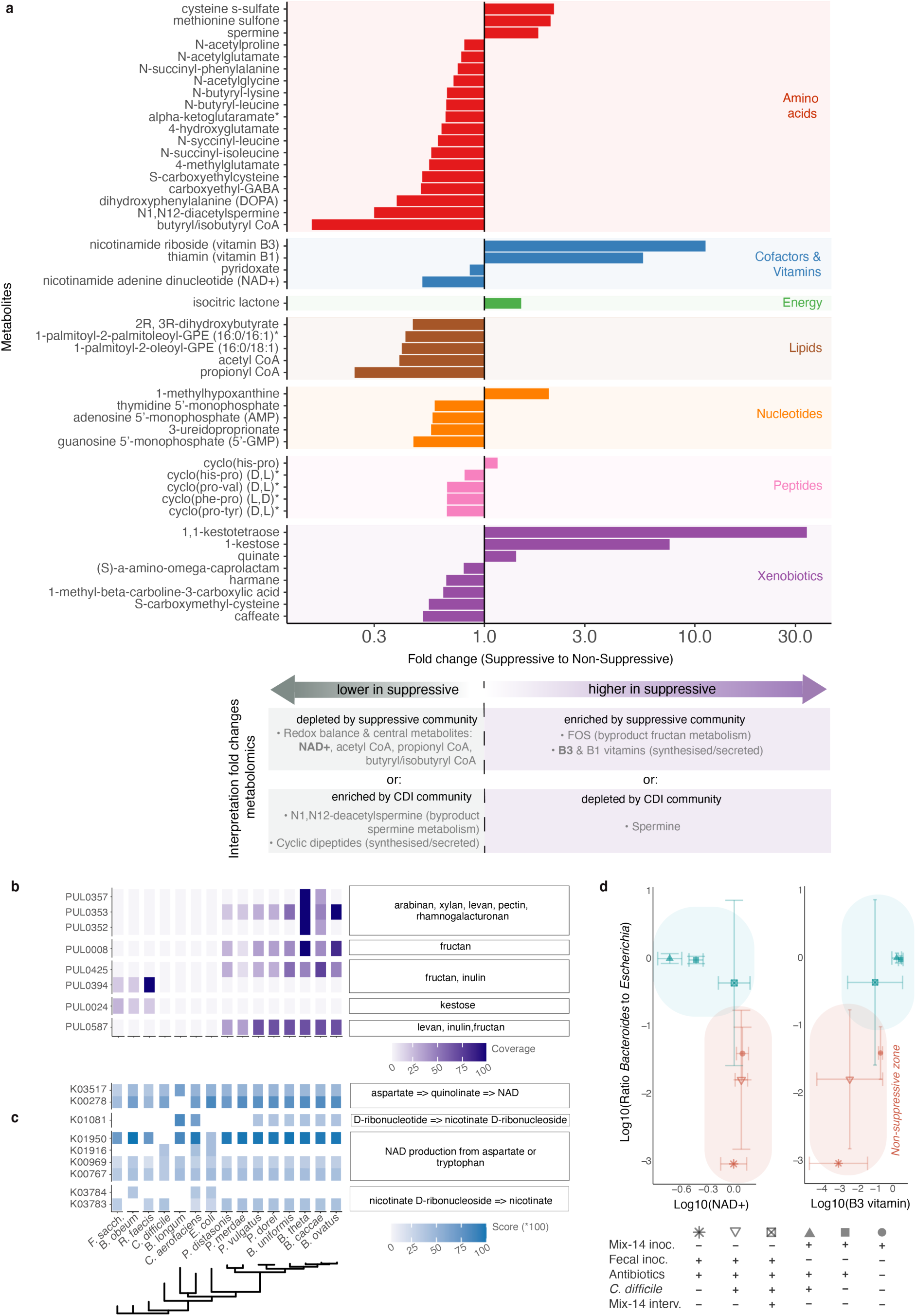
Metabolites underlying *C. difficile* suppression shared by Mix-14 and fecal-matter derived communities. **a.** Fold changes for 47 compounds identified as significantly altered (Mann-Whitney U, BH-adjusted p-value <0.05) in *C. difficile*-suppressive versus non-suppressive communities of Mix-14 and fecal origin. Compounds higher in suppressive samples have fold changes greater than 1, indicating enrichment, while those lower have fold changes less than 1, indicating depletion. Interpretations of these changes are summarized in the legend below the graph. **b.** Heatmap displaying fructan degradation gene hits from the dbCAN-PUL database across Mix-14 member and *C. difficile* genomes. Mix-14 species with the (predicted) ability to degrade the fructooligosaccharide (FOS) kestose were those that were inhibited by antibiotics and/or *C. difficile* (i.e., *F. saccharivorans, R. faecis, B. obeum*), while species that were boosted in these conditions (i.e., *Bacteroides/Phocaeicola* species) had hits in genes that could metabolise fructan into FOS such as kestose. Combined, these findings could help explain the observed spike in FOS as shown in **a.** Species were ordered by phylogenetic similarity. **c.** Heatmap displaying B3 *de novo* biosynthesis gene hits from the diamond sequence aligner using a targeted database (Methods) across Mix-14 member and *C. difficile* genomes. While most contained hits across (parts of the) the B3 biosynthesis pathway, only *Bacteroides/Phocaeicola* species contained hits across all four mapped pathways. Species were ordered by phylogenetic similarity. **d.** Ratio (log-transformed) of *Bacteroides/Phocaeicola* to *Escherichia* species consistent with differential abundance of metabolites in suppressive versus non-suppressive samples. Here shown are NAD+ (depleted in suppressive samples, showing a negative relationship with the ratio) and NAD+ precursor B3 (enriched in suppressive samples, showing a positive relationship with the ratio). An increase in vitamin B3 in conjunction with NAD+ depletion may indicate that *C. difficile* was unable to sufficiently use B3 to replenish the NAD+ required for its growth via e.g., Stickland metabolism. Thus, the suppressive community and its key interactions, including those between *Bacteroides* and *Escherichia*, may have suppressed *C. difficile* by monopolizing this vitamin and disrupting the redox balance necessary for *C. difficile*’s growth, and/or by competing with *C. difficile* for Stickland precursors such as tryptophan. Error bars represent ± standard deviations away from the log10-transformed mean ratio for y-axis, and log10- abundance of either metabolite on the x-axis.

Two of the top metabolites enriched in suppressive communities were the fructooligosaccharides (FOS) 1,1-kestotetraose (fold change = 34.0, adjusted p-value = 0.034) and 1-kestose (fold change = 7.6, adjusted p-value = 0.048). This is in line with nondigestible oligosaccharides enhancing colonization resistance against *C. difficile in vitro* [43]. FOS are known byproducts of fructan metabolism, including by *Bacteroides thetaiotaomicron:* one of the members of Mix-14’s suppressive subcommunity [44]. Inulin, a fructan, was a major component of the growth medium, and thus a likely source of 1-kestose and 1,1-kestotetraose catabolism. The increased abundance of these two compounds, coupled with the observed evidence of widespread fructan (e.g., levan, inulin) metabolism in the genomes of Mix-14 members (Fig. 4b), suggests that fructan metabolism and associated increase in fructooligosaccharides are important drivers of *C. difficile* suppression.

Nicotinamide riboside, vitamin B3, is another highly enriched metabolite (fold change=11.2, p-adjusted=0.00016). Vitamin B3 is a NAD+ precursor that can be synthesized *de novo* by *Bacteroides* [12] [45]. Enrichment of this metabolite in suppressive samples suggests a potential shift toward *de novo* B3 biosynthesis, which could result in competition with *C. difficile* for amino acid precursors like tryptophan and may alter the availability of NAD+ [46]. Genes in vitamins B1 and B3 biosynthesis pathways were detected in genomes of Mix-14 *Bacteroides/Phocaeicola* (Fig. 4c): a key genus for suppression in Mix-14 alone as well as fecal backgrounds. These metabolic shifts are associated with the changes in *Bacteroides* to *Escherichia* ratio (Fig. 4d) supporting the functional role of vitamin biosynthesis in the pathogen suppression.

Three of the top metabolites depleted in suppressive samples compared to non-suppressive were the short chain fatty acyl-CoAs: butyryl/isobutyryl CoA (fold change = 0.15, adjusted p-value = 0.00032), propionyl CoA (fold change = 0.24, adjusted p-value = 0.00074), and acetyl CoA (fold change = 0.39, adjusted p-value = 0.023). The polyamine N1,N12-diacetylspermine (fold change = 0.3, adjusted p-value = 0.037) was similarly enriched in non-suppressive samples, hinting at *C. difficile’s* growth via polyamine (i.e., spermine) metabolism. Finally, NAD+ (fold change = 0.5, adjusted p-value = 0.0014) was depleted in suppressive samples relative to non-suppressive. The enrichment of NAD+ and short chain fatty acyl-CoAs, which are involved in the regeneration of NAD+ [47, 48], in non-suppressive samples points towards their importance in redox homeostasis for *C. difficile*’s growth. Moreover, the Wood-Ljungdahl pathway has been shown to be upregulated during pathogenesis [49] and results in the production of acetyl-CoA and NAD+. These results are concordant with *C. difficile* inducing NAD+ regenerating pathways in response to availability of microbiota-produced metabolites following antibiotics treatment [50]. Moreover, the availability of NAD+ (relative to NADH) was demonstrated to play a significant role in the regulation of toxin production [51].

Cyclic dipeptides were also enriched in non-suppressive samples (mean fold change = 0.7, adjusted p-value < 0.001). *C. difficile* was previously shown to release proline-based cyclic dipeptides, D/L cylco(Leu-Pro) and D/L cylco(Phe-Pro), during colonization and under antibiotic-induced conditions [52]. These molecules were shown to be quorum sensing as well as having antibacterial properties which have been proposed to be involved in *C. difficile* colonization and pathogenesis [52, 53]. Accordingly, we observed four proline-based cyclic dipeptides enriched in non-suppressive samples: cyclo(Pro-Tyr) (D,L), cyclo(Phe-Pro) (D,L), cyclo(Pro-Val) (D,L), and cyclo(His-Pro) (D,L), suggesting that they may be a strategy employed by *C. difficile* during invasion. Furthermore, cyclo(Pro-Val) (D,L) was shown to be a diketopiperazine and beta-glucosidase inhibitor [54], suggesting a mechanism by which inulin degradation can be modulated by *C. difficile*.

### Fructan metabolism is associated with *C. difficile* suppression

Mechanistically, fructooligosaccharides likely inhibit *C. difficile*’s growth through decreasing its adhesion capacity and thus making it more susceptible to wash-out [55]. Longer fructopolysaccharides like inulin do not impact adhesion [55], thus implying that microbial community metabolism is essential. To identify which species were responsible for the observed fructooligosaccharides (i.e. kestose and kestotetraose) peak in suppressive samples (Fig. 4a), we searched the corresponding genomes for the genes known to be involved in fructan metabolism. Although some bacteria have been shown to biosynthesize fructooligosaccharides *de novo*, there were no associated hits for biosynthesis across genomes of Mix-14 members. On the other hand, 211 hits were associated with 14 fructan degradation polysaccharide utilization loci (PULs). Notably, *B. longum* and *C. difficile* lacked any fructan degrading PULs. Yet, *B. thetaiotaomicron* (SEC-species), *B. ovatus*, *B. caccae*, and *B. uniformis* possessed 50, 36, 26, and 22 fructan degradation PULs, respectively (Fig. 4b). Of all Mix-14 species, *F. saccharivorans* had the most genes annotated for the metabolism of kestose (PUL0024), contrasting with most species that had very few to none, like *E. coli* or *C. aerofaciens* possessing one and two hits, respectively.

Several species that performed well in communities after antibiotics treatment or *C. difficile* challenge, including key player *Bacteroides caccae*, thus encode genomic capabilities to metabolise fructan sources into fructooligosaccharides, while species that can metabolise kestose, such as *F. saccharivorans* and *R. faecis*, were sensitive to either perturbation (Fig. 2c, Fig. 4b, Suppl. fig. 4b). The observed fructooligosaccharides increase thus likely results from increased fructan metabolism and reduced kestose metabolism (Fig. 4a). In addition, a recent study emphasised how *Bifidobacterium* and *F. saccharivorans* abundance markedly increased in human feces following 1-kestose treatment, suggesting benefit or growth on this fructooligosaccharide *in vivo* as well [56]. Further, a recent, large-scale gut study provides evidence of *Bifidobacteria*’s fructooligosacharide utilization [57]. Moreover, [58] demonstrated that successful FMT treatment (and thus *C. difficile* suppression) in pediatric recurrent CDI (rCDI) patients was linked to consistent compositional shifts – including an increase in *Bacteroidaceae* and decrease in *Enterobacteriaceae* – correlating with an enhanced capacity for complex carbohydrate degradation. These observations further support the translational nature of the link between *C. difficile* suppression, ratio of *Bacteroides* to *Escherichia*, and complex sugar metabolism as found in our study.

### *Bacteroides* to *Escherichia* ratio, fructan metabolism, and Stickland metabolism are linked with *C. difficile* suppression in patient cohort data

To confirm the translational relevance of the compositional and metabolic signatures of *C. difficile* suppression observed in our bioreactor experiments, we analyzed 689 metagenome fecal samples from 10 published *C. difficile* cohort studies (Fig. 1a, Methods). To do so, we checked for the abundance of key genes involved in the fructooligosaccharides (FOS), Stickland, and B3 metabolic pathways among metagenomes of healthy, FMT-treated, and *C. difficile* infected (CDI) individuals (Fig. 1a, Suppl. Table 1). We found significantly fewer gene hits for all three pathways across non-suppressive (CDI) fecal samples compared to suppressive ones (healthy, post-FMT) (Fig. 5a). This result indicates these metabolic pathways might have a role in suppressing CDI *in vivo*.

**Fig. 5.**
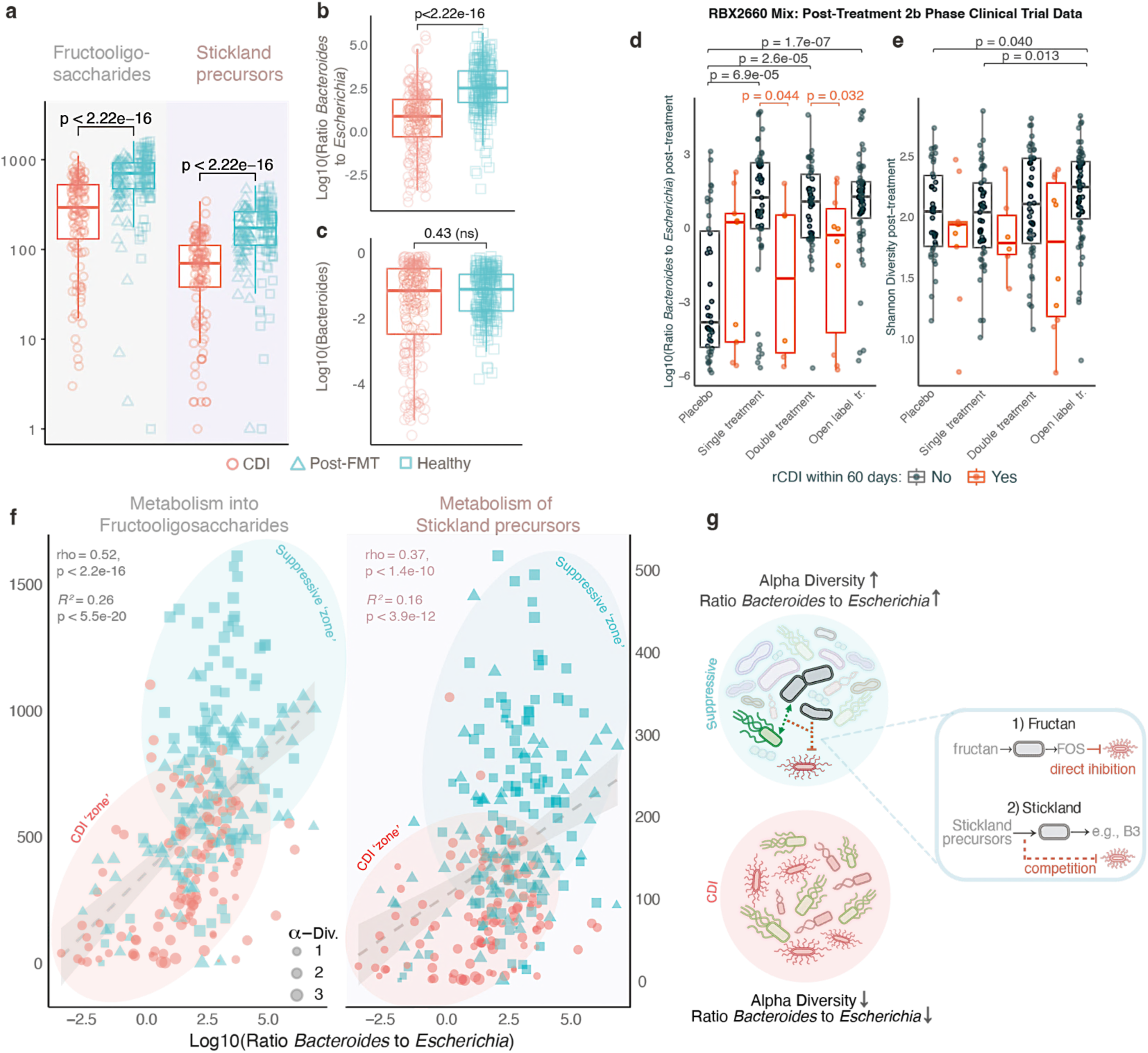
Functional and compositional signatures of *C. difficile* suppression in cohort studies support in vitro findings and mathematical modelling. **a.** FOS and Stickland precursor gene hits found in the metagenomic datasets of the analyzed cohort studies (Methods). Metabolomics of Mix-14 and fecal community supernatants indicated both FOS-generating and Stickland precursor metabolism as potential mechanistic pathways driving the observed *Bacteroides (caccae)*-mediated suppression *in vitro*. Here, we observe that genes embedded within either of these metabolic pathways are indeed found in significantly higher numbers (Mann-Whitney U tests) in fecal samples of suppressive (Healthy + FMT) than non-suppressive (CDI) human hosts, validating their potential mechanistic role in suppression. Boxplot center lines represent respective medians, box limits represent upper and lower quartiles, and whiskers represent 1.5x interquartile range. **b-c.** While the ratio of *Bacteroides* (including *Phocaeicola*) to *Escherichia* is significantly higher in healthy human samples than in CDI patient samples (**b)**, the relative abundance of *Bacteroides* is not significantly different (**c**) - consistent with *in vitro* findings (Fig. 2 e). This consistency between *in vivo* and *in vitro* data emphasizes the importance of inspecting ecological networks/configurations as potential biomarkers of *C. difficile* suppression. Boxplot center lines represent respective medians, box limits represent upper and lower quartiles, and whiskers represent 1.5x interquartile range. **d-e.** RBX2660 probiotic mix for standardized live bacterial suspension was used to treat CDI patients against rCDI with a high success rate [59]. We here investigated the ratio of *Bacteroides* to *Escherichia* (**d**) and Shannon alpha diversity calculated over Genus-level (**e**) between the different treatment groups. For post-treatment days datapoints (Day >0), statistical comparisons (Mann-Whitney U test, BHadjustments) were performed between treatment types (black brackets, shown if p < 0.05) and within treatment types (rCDI within 60 days, red brackets, shown if p < 0.05). Open label treatments were excluded from within-treatment comparison, since all patients had rCDI prior to receiving the open label treatment. Between treatment types, ratio is a better discriminator between placebo and non-placebo treatments, successfully distinguishing all treatment types (single-, double-, and open label treatments) from placebo (p < 0.05), while alpha diversity only successfully distinguishes placebo from open label treatment patients, and shows significant differences within treatments (e.g., single treatment versus open label treatment). Within treatment types, ratio is a better discriminator between successful and unsuccessful patients, showing a significantly lower ratio in patients for whom the single or double dose of did not successfully prevent rCDI within 60 days. Alpha diversity failed to have the same discriminatory power – only the comparison between successful and unsuccessful patients for double dose reaching near statistical significance (p = 0.054). Boxplot center lines represent respective medians, box limits represent upper and lower quartiles, and whiskers represent 1.5x interquartile range. **f.** Both the ratio of *Bacteroides* to *Escherichia* and hypothesised metabolic gene hits were found to be significantly different between fecal samples of suppressive and non-suppressive hosts (see panel **a-c**). Here, we show that these compositional and molecular signatures are positively related, as were predicted from our *in vitro* data (Fig. 4). The fitted line represents the linear regression, and the shaded area the associated standard error. **g.** Proposed phenomenological as well as mechanistic signatures of *C. difficile* suppression: non-suppressive communities are defined by lower alpha diversity and lower ratio of *Bacteroides* (incl. *Phocaeicola*) to *Escherichia*, while suppressive communities share compositional traits, including higher alpha diversity and higher ratio of *Bacteroides* to *Escherichia*. These traits were statistically linked to two potential molecular mechanisms: 1) fructan metabolism into FOS, such as 1-kestose, which may directly suppress *C. difficile,* and 2) metabolism of Stickland precursors – including NAD+ and cofactors, as well as tryptophan for *de novo* B3 biosynthesis – indirectly suppresses *C. difficile* via competition for critical substrates and redox balance/central metabolites.

Furthermore, we checked whether the difference in bacterial ratio (*Bacteroides* to *Escherichia*) observed in our *in vitro* studies was also present in patient metagenomic samples. As hypothesized, the *Bacteroides* to *Escherichia* ratio was significantly lower in the CDI patients compared to the healthy individuals (Fig. 5b). Since *Bacteroides* relative abundance was not significantly different between the suppressive and non-suppressive communities (Fig. 5c), this confirms the importance of the ratio rather than *Bacteroides* alone in the association with suppression. To further assess whether the ratio of *Bacteroides* to *Escherichia* might serve as an important variable in explaining or predicting the success of designed bacteriotherapeutic interventions for treating recurrent CDI, we also analyzed the metagenomic datasets of a study [59] showing the potential of a live biotherapeutic mix (i.e., “second generation FMT”), RBX2660, that was shown to be effective in recovering patients from rCDI (Suppl. fig. 14). The dataset includes a subset of cases of patients who were treated with the mix but still suffered from rCDI within 60 days post-treatment. We observed that the ratio of *Bacteroides* to *Escherichia* was significantly lower (Mann-Whitney U test, BH-adjusted p < 0.05) both between placebo and treated patients, as well as successfully treated and unsuccessfully treated patients (Fig. 5d, Suppl. fig. 13). While the *Bacteroides* to *Escherichia* ratio strongly distinguished between placebo and all treatment groups (i.e., single-, double-, or open-label treatments) at post-treatment stages (p < 3e-05, Fig. 5d & Suppl. fig. 13), Shannon alpha diversity is a less effective differentiator; within treatment groups, it did not statistically differentiate successful from unsuccessful treatments. Between treatment groups, the only significant differences in alpha diversity were between open-label treatment and placebo (p = 0.04), as well as between single treatment and open-label treatment (Fig. 5e). This implies that alpha diversity is not a strong metric in differentiating treatment groups from placebo groups, or in differentiating successful from unsuccessful patients. Even when single dose and double dose treatment data were grouped together, these trends held (Suppl. fig. 13).

Since both the differential microbial ratio (found in the 16S data of our Mix-14 and fecal community experiments) and differential gene presence (of pathways associated with metabolites found to be differentially abundant in our experiment’s metabolomics data) were found in the patient metagenome data, we further tested whether these signatures were related. As shown in Fig. 5f, the *Bacteroides* to *Escherichia* ratio positively correlated with the abundance of fructooligosaccharide- and Stickland metabolism related genes, revealing clusters of CDI (low alpha diversity, low gene counts and low ratio) and suppressive (high alpha diversity, high gene counts and high ratio) patient fecal communities (Fig. 5f). Similar clustering was observed for vitamin B3 biosynthesis pathway-related genes (Suppl. fig. 15). Altogether, our results indicate that suppressive communities (healthy and FMT-treated) show an increase in associated *Bacteroides* to *Escherichia* ratio, which in turn is associated with fructooligosaccharide- and Stickland metabolism pathways. Finally, our findings from *in vitro* bioreactor-based experiments with mathematical modeling were consistent with patient metagenome samples, showing the *in vivo* relevance of the methods utilized.

## DISCUSSION

The suppression of enteric pathogens such as *C. difficile* by gut microbiota in controlled by complex and multifactorial mechanisms [60, 61], including direct inhibition [62], enhancement of the host immune response [63, 64], nutrient competition [65, 66], and modulation of secondary metabolism [67]. The success of FMT in suppressing CDI suggests that multiple mechanisms need to be operating simultaneously to achieve pathogen suppression. Some of these mechanisms are well understood, such as the inhibition of *C. difficile* associated with the conversion of primary bile acids into secondary bile acids [68, 69] and the increased production of short-chain fatty acids [70]. However, the overall outcome of pathogen suppression can vary with subtle changes in microbial community interactions. For instance, some cases of FMT still fail to resolve rCDI [71]. A deeper understanding of microbial interactions is therefore necessary to develop defined bacterial therapeutics, improve treatment outcomes, and to identify microbiome-based biomarkers for stratifying treatment strategies.

Together, our experiments using synthetic and fecal-matter-derived communities, metabolomics data, mathematical modelling, and cohort metagenomics data analysis uncover the ratio between *Bacteroides* and *Escherichia* as a key feature of *C. difficile* suppressive capacity of a gut microbial community. While this implies that having no *Escherichia* would impart higher colonization resistance, mathematical modelling of our synthetic gut community suggests that relatively low levels of *E. coli* contribute to community suppressiveness by positively interacting with *B. caccae*. This non-intuitive conjecture of the model was validated *in vitro*, wherein a combination with *E. coli* showed higher suppressive capacities compared with *B. caccae* or other suppressive species alone.

Notably, *E. coli*’s support of *B. caccae* is key, further enabling commensal or mutualistic interactions with other co-suppressors like *P. distasonis*. In the absence of *B. caccae* and other co-suppressors from the community, *E. coli* and *C. difficile* can co-exist in high relative abundances. These findings showcase the importance of non-linear, higher-order, interactions in successful pathogen suppression, and provide mechanistic insights into high variability observed across individuals and treatments, including the inconsistent success rates of bacteriotherapy in preventing or recovering recipients from CDI [72]. In addition, emergent interactions explain inconsistencies in associations between individual taxa and disease states. For instance, *E. coli* has been linked to both CDI suppression [73] and occurrence [74, 75]; and *Bacteroides –* a key co-suppressing taxon in our study – has previously been linked to CDI suppression [76] as well as CDI occurrence [2]. Pathogen suppression is thus an emergent property of community interactions.

The importance of shifting the focus away from individual taxa and towards ecological networks to link gut composition with health outcomes has also been emphasized previously in chronic fatigue [77], engraftment success [78]; and response to cancer immunotherapy [79]. The non-linear ratio signature that we observed surpasses any single taxon in characterizing successful CDI suppression, but also community-scale metrics such as alpha diversity. Community alpha diversity is currently considered a major contributor in pathogen suppression as well as many other microbiota-linked host-benefits [2, 80]. A combination of *in vitro* and *in vivo* analyses as demonstrated in this study could therefore be used to identify ecological processes governing structure-function relationship in other microbiome-linked disease states.

At molecular level, we find that the suppression of *C. difficile* is, at least partially, mediated by small molecules produced and exchanged by the suppressive sub-community. The corresponding interactions encompass both inhibition and nutrient competition with the pathogen. The inhibitory molecules identified in our study include fructooligosaccharides (FOS) like 1-kestose and 1,1-kestotetraose, which have previously been shown to modulate *C. difficile*’s adhesive phenotype [43, 44], and thereby reducing its residence time. The presence of these two compounds in the metabolomics data, coupled with the observed evidence of widespread fructan (e.g., levan, inulin) metabolism across members of Mix-14 genomes (Fig. 4b), suggests that fructan metabolism and associated increase in fructooligosaccharides is an important driver of *C. difficile* suppression.

On nutrient competition, > 40% of metabolites that were significantly lower in suppressive exometabolomes were amino acids or derivatives in accord with *C. difficile’s* dependence on Stickland metabolism [42], and showcased an enrichment of polyamine endproducts such as N1, N12-deacetylspermine in CDI communities. Similarly, other precursors and co-factors needed for *C. difficile* growth as direct substrates or to maintain redox balance, including short-chain fatty acyl-CoAs and NAD+, were also depleted in the suppressive community, redirecting these precursors and co-factors to B1 and B3 vitamins, among others. A converted form of vitamin B3, nicotinamide mononucleoside, was previously shown to be key in colonization resistance against other gut pathogens [81].

Supporting our metabolomics findings, analysis of microbiome samples from independent, previously published, cohort and clinical trial datasets identified *Bacteroides* to *Escherichia* ratio as a key compositional signature of the pathogen suppression and treatment success. We note that the analysis does not necessarily imply that the absence of *Escherichia*, which will make the ratio infinity, would be beneficial. As our modeling and in vitro experiments suggest, *Escherichia* do have a role to play in the pathogen suppression. As most, if not all, gut microbiomes harbor *Escherichia*, the *Bacteroides* to *Escherichia* ratio serves as a good indicator. Gene-level analysis of the microbiome samples further attested our mechanistic findings on the role of fructan and Stickland metabolism in *C. difficile* suppression. While the failure of FMT and other bacteriotherapies may be influenced by other recipient-specific factors, e.g. use of antibiotics [58, 82, 83], the strong cross-cohort signatures observed in our analysis and the underlying mechanistic reasoning suggests that part of the inter-individual variation in treatment response could be explained by the ecological and metabolic signatures identified in this study.

Together, our results bring forward ecological and metabolic signatures for understanding and predicting the success of bacteriotherapies and FMT in treating CDI and preventing recurrence. This represents a mechanistic and robust approach compared to low-resolution metrics like alpha diversity, which is currently thought to be the main factor underlying FMT success [84]. Our findings could be used to characterize FMT- donors and to match bacterial therapeutics to recipients using, e.g. the *Bacteroides* to *Escherichia* ratios, while measurements of metabolites like fructooliogosaccharides and vitamin B3 could be used as biomarkers to monitor treatment success.

## METHODS

### Bacterial growth conditions

All the anaerobic bacteria for the study were grown in modified Brain Heart Infusion (mBHI) broth, as in [31]. Briefly, the growth medium was prepared by adding yeast extract (5.0 g/L), L-cysteine (0.3 g/L), 1 mL/L Resazurine (0.25 mg/mL), and 0.05 g/L of bovine bile to the standard BHI base ingredients. The media was purged until anaerobic and was autoclaved at 121 degrees Celsius and 15 lbs pressure for 30 minutes. The media was then transferred to the anaerobic chamber and further supplemented with 1 mL/L of hemin solution (0.5mg/mL), 1 mL/L Menadione (5.8 mM), 10 mL of ATCC vitamin mixture (ATCC, USA), 10 mL of ATCC mineral mixture (ATCC, USA), 8mL/L of 10N NaOH, and 100 mL/L of 1M 2-(N-Morpholino) ethane sulfonic acid (MES) (Table 1). For making *C. difficile* selective agar, mBHI was supplemented with 0.3g/L of D-cycloserine and 0.002g/L of Cefoxitin. Medium composition and procedure for making the dGMM+Lab media were taken from [85].

### Fecal samples preparation and mini bioreactor

Donor fecal samples were collected as described in [31]. Briefly, we obtained fresh fecal samples from six healthy individuals with no record of antibiotic use in the past three years. The six fecal samples were pooled equally and cryopreserved at −80°C with 10% DMSO as a cryoprotectant, and used as inoculum. A similar approach was used to prepare Mix-14, Mix-4, and Mix-3 stocks (Fig. 1c,d). All the individual bacteria were grown until they reached an optical density of 0.5 or diluted to it, if necessary, with mBHI and then cryopreserved in −80°C with 10% DMSO as a cryoprotectant, and were subsequently used as inoculum. mBHI medium was used as a culture medium for the complete bioreactor run. Mini bioreactors (MBRAs) were sterilized and assembled, after which the experiment was performed as described previously in [40], with minor modifications. The input and output on Watson Marlow pumps were set at 1 rpm and 2 rpm, respectively. The rotating magnetic stirrer was set at 150 rpm with a 10mm stir bar inside the bioreactor wells. Four reactor blocks were used for every run, containing six wells per block, producing a total of 24 wells per run. Per run, the wells were divided into 4-5 treatment groups. Each treatment group was thus allocated 4 or 6 replicate wells. The growth medium was set to flow continuously into all reactor wells, consuming ∼300mL for 24 hours. Per run, 300μL of inoculum (for Mix-14 or fecal community) was introduced into all wells, during which the flow was stopped for 16 hours to allow engraftment and reduce the role of bottlenecking during community assembly. Upon flow resumption, the continuous flow model was operated up to 23 days post-inoculation. On day 8, growth medium containing a concentration of 25 mg/L of clindamycin was used for respective treatment conditions (Fig 1c,d), and discontinued on the 12^th^ day of the reactor run, after which flow with regular growth medium was resumed. On the 13^th^ day of the reactor run, 150 μL of *Clostridium difficile* R20291(OD 0.16) was inoculated into all the reactor wells of respective treatment conditions. On day 16 of the run, we inoculated wells of respective experimental conditions with 300μL (OD 0.5) of Mix-14, Mix-4, Mix-3, or individual bacteria (Fig. 1d). Samples were collected daily from each reactor well to determine 16S rRNA analysis, metabolomics, OD at 600nm, and pH.

### DNA isolation and sequencing

For Mix-14 experiments (Fig 1c), DNA was isolated from all samples extracted daily from respective bioreactor runs. For the experiment with fecal-matter derived communities (Fecal + *C. difficile*, Fecal + antibiotics + *C. difficile*, Fecal + antibiotics + *C. difficile* + Mix-14), DNA was isolated from the initial inoculum (day 0), day 8, day16, day 20 and day 23 (Fig. 1d). For follow-up experiments with the fecal community (e.g., intervention by individual bacteria, Fig. 1d), DNA was isolated from day 20 samples. DNA was extracted using a Powersoil DNA isolation kit (MoBio Laboratories Inc, CA) following the manufacturer’s instructions. After extraction, the quality of DNA was measured using NanoDropTM one (Thermo Fisher Scientific, DE) and quantified using Qubit Fluorometer 3.0 (Invitrogen, CA). The DNA samples were stored at −20°C until further use. To examine the change in microbial composition over time, all samples were amplicon sequenced using an Illumina MiSeq platform with paired-end V3 chemistry. The library was prepared using an Illumina Nextera XT library preparation kit (Illumina Inc, CA) targeting V3-V4 regions of the 16S rRNA sequence. The libraries were bead normalized and multiplexed before loading into the sequencer.

### 16S Amplicon sequencing analysis

Microbiota profiling from 16S sequencing was performed using Vsearch [86]. Merging and Quality filtering with a minimum cut-off length of 400 and max length of 500 was done for the fastq files using the Vsearch tool. Singleton and chimeric reads (UCHIME) were removed. OTU picking was performed with VSEARCH abundance-based greedy clustering. OTUs were annotated with the SILVA reference database or custom reference database consisting of full-length 16S extracted from the whole genome sequences of the fourteen bacteria in the consortium.

### *In vitro* paired co-culture inhibition (relevant for Fig. 1b)

Co-culture inhibition was performed as previously mentioned in [31] with slight modifications. Briefly, Mix- 14 member species and *C. difficile* were grown in mBHI medium overnight until they reached an OD (600nm) of 0.8-1. Each culture was then diluted to 0.5 OD using an mBHI medium. For the paired co-culture assay, different ratios of Commensal:Pathogen (1:1, 1:4, and 1:9) were diluted in 1mL of fresh mBHI or dGMM&LAB medium and incubated for 24 hours. Each culture was then plated on CDSA plates at different dilutions and incubated for 24 hours. Colony counts were done, and percentage inhibition was subsequently calculated in units of CFU/mL *C. difficile*.

### Antimicrobial inhibition (relevant for Suppl. fig. 12)

For the antimicrobial inhibition assay, *B. caccae* strains were grown for 24 hours in mBHI medium and then centrifuged at 1000 g for 5 min. The supernatant was filter-sterilized using a 0.22 μm filter in the anaerobic chamber and diluted to a ratio of 1:1 with the 1X mBHI medium. pH was adjusted to 6.8. An overnight culture of *C. difficile* was diluted to OD600 of 0.5. Twenty microliters of OD600-adjusted suspension was added to 1 mL of the 1:1 diluted cell-free supernatant and incubated for 24 hours in triplicates. *C. difficile* was grown in 50% mBHI diluted with anaerobic PBS as a positive control. After 24 hours, the cultures were serial-diluted with anaerobic PBS, plated on CDSA, and incubated anaerobically for 24 hours for enumeration. For heat treatment, the centrifuged supernatant was transferred to sterile tubes and pasteurized at 90 °C for 45 min (1 mL portions) suspended in a water-filled heating block. Pasteurized supernatant preparations were stored at 4°C for near-term experiments or frozen at −20 °C. For Proteinase K, portions of supernatants were treated with proteinase K at 1mg/ mL concentration for 1 hour at 37°C. Following treatment, the enzyme was inactivated by the addition of phenylmethylsulfonyl fluoride.

### *In vitro* growth curve analysis (relevant for Suppl. fig. 2)

For growth curve analysis, individual bacteria were grown in the mBHI medium until reaching the mid-log phase at OD 0.8-1. Then, each monoculture was diluted to an OD (600nm) of 0.005, and OD at 600nm was measured regularly for 96 hours, allowing all bacteria to reach the stationary phase. All the bacteria were grown in biological triplicates.

### Community-level physiology profiling and Biolog metabolite utilization profiling (relevant for Suppl. fig. 10)

Biolog data for assaying carbon utilisation profiles of individual bacteria was extracted from previous work [31] [87]. For community-level physiology profiling, 1mL samples were collected from the bioreactor run of different treatments. The samples were centrifuged at 3000 x g for 1 min. The pellets were washed thrice with anaerobic PBS until the residual media was thoroughly washed off. The remaining pellet was dissolved in enough AN inoculating fluid (Biolog Inc.), to reach a final OD (600 nm) of 0.02. The AN fluid containing bacterial cells was then aliquoted 300μL to individual wells in the AN Biolog plate. OD at 600 nm was measured to verify the OD and further checked at 12 and 24 hours. For the analysis, a cutoff of 20% utilization was set to determine the positive utilization of the substrate.

### Semi batch *in vitro* coculture experiment (relevant for Fig. 3c)

We did an *in vitro* semi-batch coculture inhibition assay to validate mathematical modeling results. A mix of different bacteria was made, as mentioned previously. 150uL of the bacterial mix was inoculated onto 10mL of mBHI. The culture was half diluted with mBHI every 24 hours for five days. 100μL of 0.19 OD(600nm) *C. difficile* was inoculated to the culture, and the CFU count was done in CDSA for 2 days post-infection. Cultures were collected for DNA isolation before and post-infection.

### Antibiotic susceptibility test (relevant for Suppl. fig. 3)

An antibiotic susceptibility test is done to determine the ability of the core human gut bacteria to resist the antibiotic used for dysbiosis. Individual bacteria were grown in the mBHI media until they grew to the mid-log phase OD 0.8-1. Then, the culture was diluted to become an OD of 0.005 and grown until it reached 0.2. Further, clindamycin was added at a concentration of 250𝜇g/mL. After 36 hours, the amount of total ATP produced was measured using a luminescence assay kit according to the manufacturer’s instructions (Promega). Fold change was calculated for the relative luminescence against non-antibiotic treated control.

### Untargeted Metabolomics (relevant for Fig. 4a)

Samples from the bioreactor run on day 16/20 were filtered through a 0.2𝜇m filter and sent to Metabolon Inc. for analysis. Samples were processed according to the protocols for metabolomic analysis provided by Metabolon Inc. Details of sample accessioning, preparation, quality control measures, and ultra-high-performance liquid chromatography-tandem mass spectroscopy (UPLC-MS/MS) can be found in the resources provided by Metabolon [88] and are included in the Supplementary Methods.

### Community dynamics data analysis (relevant for Fig. 2)

For experiments with the defined Mix-14 community, 16S amplicon sequencing reads were normalized to arrive at relative abundances for each member species across all six replicates for each experimental condition. Statistical comparisons of endpoint relative abundance profiles (day 23 for ‘Mix-14 control’, ‘Mix-14 + Antibiotics’, ‘Mix-14 + Antibiotics + *C. difficile*’, and day 14 for ‘Mix-14 + *C. difficile’*) were performed using the Mann-Whitney U test, as implemented in the ‘geom_signif’ function of the ‘ggsignif’ R package (v 0.6.4).

To construct correlation matrices for Mix-14 relative abundance profiles, we merged all endpoint relative abundance data across all four experimental conditions. Binary variables were added to account for absence or presence of antibiotics treatment or *C. difficile* inoculation. Spearman correlations between all fourteen Mix-14 member species and their end-point relative abundances, as well as the two binary experimental perturbation variables, were calculated, arriving at a 16 x 16 correlation matrix using the ‘rcorr’ function of the ‘Hmisc’ R package (v 5.1-0). Correlation networks of significant (p<0.05) correlations were constructed using the ‘igraph’ R package (v 1.4.2).

In experiments in which an undefined community was assembled from a fecal background, the maximum resolution of taxonomic assignment from 16S amplicon sequencing was genus level. Relative abundances were calculated for each assigned OTU, and those with the same genus name were summed together. Since *Escherichia* was indistinguishable from *Shigella,* they were grouped together in their OTU assignments. Endpoint (day 20) relative abundance profiles for all ten fecal background community experiments were assessed together and assigned a binary value of suppressiveness of *C. difficile* (1 for log10 CFU/mL counts of < 2, and 0 for > 2).

To calculate the Bray-Curtis dissimilarity of final mean relative abundance profiles across the different experimental conditions for both Mix-14 as well as Fecal background communities, we used the ‘vegdist’ function of the ‘vegan’ R package (v 2.6-4). To plot the Bray-Curtis dissimilarities in a heatmap, we used the ‘pheatmap’ function of the ‘pheatmap’ R package (v 1.0.12).

For all endpoint relative abundance profiles (of Mix-14 defined and fecal undefined communities), we calculated the corresponding Shannon alpha diversity index (referred to as alpha diversity or Shannon Index) and the 1:1 (56 in total) and 2:1 ratios (168 in total) of Mix-14 genera. Shannon alpha diversity index was calculated as follows: 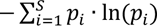, where 𝑆 is the total number of taxa (with nonzero relative abundance), and 𝑝 is the relative abundance of taxon 𝑖. The 1:1 ratios were calculated via 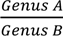, while the 2:1 ratios were calculated via 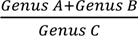. Prior to calculating ratios, a small (0.0001) value was added to all relative abundance data so that ratio calculations would not result in 0 or ‘inf’ values, which in turn would create issues when log10-transforming ratios. Log10-transformation of the ratios was performed to account for the wide range of values and to normalize the distribution, making it more suitable for statistical analysis and easier to interpret visually on graphs.

Statistical significance of difference in Shannon index between unperturbed versus perturbed Mix-14 and non-suppressive versus suppressive fecal communities was calculated using the Mann-Whitney U test (pairwise Wilcoxon rank sum test), implemented in the ‘wilcox.test’ function of the default ‘stats’ R package (v 4.2.2). The same analysis was done for 1:1 and 2:1 log-10 transformed ratios, where effect sizes were calculated by taking the difference in medians for each ratio between compared groups. To emphasize potential ecological configurations of relevance in suppressive mechanisms in both Mix-14 as well as fecal background experiments, ratios that were significantly different in both comparisons of perturbed versus unperturbed Mix-14 and suppressive versus non-suppressive fecal communities, while sharing a similar effect direction, were selected for further analysis.

To understand the relationship between fecal background community suppressiveness, alpha diversity and selected ratios, we executed linear regressions on log-transformed ratios against corresponding alpha diversity. The linear regressions (and *R^2^* calculations) were performed using R’s baseline ‘lm’ function, and the statistical significance of *R^2^* was assessed using the function’s default F-test.

### Ecological Modelling (relevant for Fig. 3b)

To explore to what extent the observed dynamics and properties of the experimental Mix-14 communities can be recreated *in silico* based on observed monoculture growth kinetics and/or suppressiveness *in vitro*, we here formulate a model mimicking the bioreactor experimental timeline and constraints. Moreover, to further study the observed emergent and key role of *B. caccae* on Mix-14’s suppressive capacity and Shannon-indexed alpha diversity, we will here focus on the following key Mix-14 clusters/players (>10% final abundance in absence or presence of antibiotics treatment, and/or those with individual suppressive capacities): *B. caccae, E. coli, F. saccharivorans*, SEC-species and *C. difficile*.

#### Model formulation and integration

A system of coupled ordinary differential equations, following the generalized Lotka-Volterra formulation, was used to investigate which interaction landscape(s) are most likely to explain Mix-14’s post-antibiotics suppressiveness of *C. difficile* (equations 2-4). Parameter inference was performed using supplied experimental data, such as monoculture growth kinetics of each Mix-14 community member.

To calculate the maximum growth rate for each species, we determined the maximum change in OD value per unit time as measured in monoculture. Given that the time between subsequent timepoints was not always equal, we used the finite difference method for approximating the derivative:

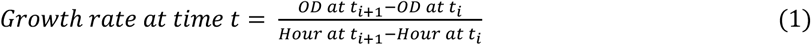

After which we extracted the mean maximum net hourly growth rate for each species. Since the unit of time used in the model is day^-1^, we multiplied the maximum net hourly growth rate by 24 to arrive at a daily maximum net growth rate. Then, the dilution rate (𝜙) was added to arrive at the per capita growth rate (since the net growth rates represent the difference between inherent growth rate and the flush effect/dilution rate of the bioreactors, which is proportional to total biomass as well). For the SEC-species (*B. dorei, B. thetaiotaomicon, P. distasonis*), the mean maximum per capita growth rates and corresponding standard deviations were extracted for all member species and averaged across them. As described further in Results (2.4., Suppl. fig. 6), the SEC-species all positively covaried in Mix-14 post-antibiotic treatment and all displayed individual suppressive capacities (Fig. 1b).

The parsimonious model system consists of the following coupled ordinary differential equations:

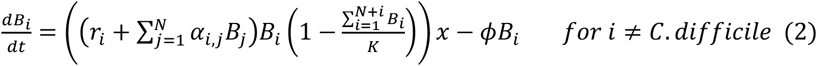

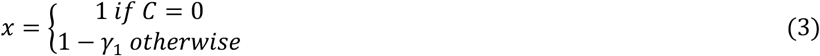

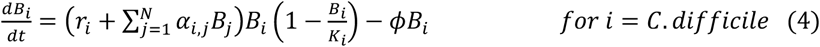

Where 𝐵_𝑖_ is the population abundance of species 𝑖, 𝑟_𝑖_ describes the per capita growth rate of species 𝑖 (informed by max growth rate in monoculture growth curves), 𝑁 describes the total number of species in the system (excluding species 𝑖), 𝛼_𝑖,j_ describes effect of species 𝑗 on the per capita growth rate of species 𝑖, 𝐾 describes the ‘carrying capacity’ or maximum density of the system (informed by range of observed co-culture max ODs), and 𝜙 describes the dilution rate of the system. The interaction coefficients were all made relative to the subjected species’ growth rate, 𝑟_𝑖_, so that one unit of the exerting species, 𝑗, corresponds to a fraction, 𝑓, of the subjected species’ growth rate being impacted, and thus 𝛼_𝑖,j_ = 𝑓 · 𝑟_𝑖_. If the direction of interaction (i.e., positive, or negative) was inferable from experimental data, the relative strengths of interaction (𝑓) were pulled from uniform distributions following 𝑓∼U(0.01, 1), otherwise: 𝑓∼N(0, 0.2^2^) (assuming neutrality, with mean of 0). A random permutation experiment (akin to [89, 90]) was thus performed on each interaction to arrive at a maximum combinatorial space, not only of interaction directions but also strengths. See Fig. 3b for configurations.

Multiplier 𝑥 describes the bacteriostatic sensitivity as a function of antibiotics treatment, with the degree of sensitivity being captured by 𝛾_1_, which is 0 for complete insensitivity, and 1 for complete sensitivity (i.e., halting growth altogether). As evident from studying relative abundance dynamics of the co-cultures in the window between antibiotics addition (day 8) and subsequent two days of treatment (day 10), apparent responses to antibiotics in community co-culture are different – or altogether opposite – to antibiotic-sensitivity measured in monoculture (Suppl. fig. 3). This hints at the importance of ecological forces such as community cross-protection or community sensitization: effects that result from interaction networks within the community, reducing the accuracy at which monoculture growth data can be extrapolated to community growth [91, 92]. In the hypothesised ecological trajectory of suppression, this community-induced sensitivity of *F. saccharivorans* is a critical point in the community’s dynamics, directly or indirectly allowing for key player *B. caccae* to take over *F. saccharivorans’* dynamic niche (i.e., as major grower alongside *E. coli,* oscillating along the system’s maximum optical density). Consistent between monoculture and co-culture is *E. coli*’s response to the antibiotics treatment, which has been shown to be bacteriostatic [93]. For modelling purposes, we here include community-level rather than monoculture sensitivity to antibiotics in multiplier 𝑥, avoiding a significant increase in parameter space (via adding an antibiotics-dependency to each interaction coefficient in the 𝛼_𝑖,j_ matrix). Moreover, since *C. difficile* is introduced days after the last antibiotic treatment, we assume antibiotics are flushed out of the system and hence do not include multiplier 𝑥 for *C. difficile*.

By enforcing a universal carrying capacity in the form of 𝐾, all subjected populations suffer equally from the substrates being utilised fast when the bioreactor reaches high population densities. The universal *K* does slow down population growth rates as cellular densities get higher, which is not to account for metabolic reprogramming as population reach stationary phase (which is what *K* typically represents in logistic growth models for serial dilutions), but rather to account for the (implicit) Monod growth kinetics, slowing down inherent growth rates as substrates → 0. In addition, this enforcement allows us to mimic the anti-correlative behaviour around system equilibrium observed for the ‘big players’ (i.e., *F. saccharivorans* and *E. coli* without antibiotics treatment, *B. caccae* and *E. coli* with antibiotics treatment) that are coupled in a dynamic that cannot be sustained if their interaction was inherently antagonistic [94].

The differential for *C. difficile* is taken separately, with its growth being limited not by the universal *K*, but rather by its species-specific 𝐾_𝑖_ (as extracted from the monoculture maximum). The rationale for this choice is twofold: firstly, if *C. difficile* were also constrained by a universal carrying capacity, any apparent suppression could result from the model artefact of carrying capacity saturation by day 13 (*C. difficile*’s inoculation day), rather than direct interactions with other community members. Secondly, given the opportunistic, pathogenic nature of *C. difficile*, it is biologically plausible to assert that this bacterium operates under different ecological principles compared to the resident community it is introduced to, disrupting any stabilising ecological forces at play within the community (such as being allowed to ‘parasitise’ on the system’s carrying capacity, by not being bound by it itself).

For each simulation, we collected:

i. All inputted parameters (𝑟_𝑖_, 𝛼_𝑖,j_, 𝐾);
ii. Final abundances (𝑂𝐷_𝑖_ at the end of day 23);
iii. Final community OD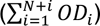 ;
iv. Final relative abundances 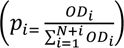;
v. End-point Shannon alpha diversity index 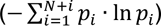
vi. Suppressiveness of final community 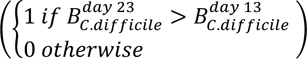
vii. Ratio of final relative abundances of *E. coli* versus *B. caccae* and SEC-species 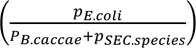.

Simulations producing a final community composition in which the following conditions were met, were kept (i.e., those representing a realistic interaction network producing observed trends for Mix-14 treated with antibiotics):

i. *B. caccae*’s final relative abundance > 0.2; and
ii. SEC-species’s summed final relative abundance > 0.02; and
iii. *E. coli* > *B. caccae;* and
iv. *E. coli* > SEC-species.

The model was integrated using the ‘ode’ function of the ‘deSolve’ R package (v 1.35). Each simulation was initialised with identical starting ODs for each state variable (0.01 OD, except *C. difficile* with an initial OD of 0), to reflect low initial densities in the order of magnitude used in the experiment and was integrated for 23 days in a chemostat environment mimicking the experimental setup. The antibiotics effect multipliers were made to apply (i.e., change from 1 to ≤1) from the start of the antibiotics treatment (day 8), until the end, including the day it takes for the antibiotics to be washed out of the system (day 12, accounting for another 3 volume changes after the end of the antibiotics treatment). Due to the above-described sources of inter-simulation variation (i.e., random permutation of interaction parameters and stochasticity on dilution rate variation), 1.5·10^6^ model integrations were conducted to extract sufficient simulations per network configuration (Fig. 3b). A final subset of 4.3·10^4^ simulations met the conditions as described above and was used for further analysis. An overview of model parameters and associated units can be found in Suppl. Table 3A.

#### Analysis simulation results

The relative importance of each interaction between the key Mix-14 players (within constraints as shown in Suppl. Table 3B & Fig. 3b) in driving system Suppressiveness and Shannon alpha diversity was investigated using the simulation output results (of 1.5·10^6^ simulations, ∼4·10^4^ met conditions as detailed above and were used for analysis). The *f* values of every interaction coefficient for each simulation were extracted, and interactions were classified as follows: 𝑓 < −0.5 = strongly negative (i.e., one unit of the effector corresponds to inhibition of at least half of the effected species’ growth rate), −0.5 < 𝑓 < −0.1 = moderately negative, −0.1 < 𝑓 < 0 = weakly negative, 0 > 𝑓 < 0.1 = weakly positive, 0.1 > 𝑓 < 0.5 = moderately positive, 𝑓 > 0.5 = strongly positive.

A binary score metric was added to the simulation results, in which 1 was assigned to simulations for which the final Shannon alpha diversity (calculated for the resident community, i.e., excluding *C. difficile*) was within the observed range of communities treated with antibiotics (i.e., Shannon index of > 0.8 but < 1.2), whereas 0 was assigned to simulation results that fell outside of the observed range. The same binary metric was added for suppressiveness, with 1 being assigned for simulations accurately reproducing *C. difficile* suppression, and 0 when a simulation result allowed for *C. difficile* growth.

Simulation results (Suppressiveness score and Shannon index range score) were binned into classifications of interaction type and strength (e.g., ‘weakly positive’ for ‘interaction effect *B. caccae* on *E. coli*), and the ratio of ‘0’ to ‘1’ scores for both the Suppressiveness score and the Shannon index range score were calculated for each bin. These binned simulation results were plotted in heatmaps using the ‘ggplot2’ R package (v 3.4.2). Ratios of <1 (i.e., 1 being an equal number of ‘realistic’ to ‘unrealistic’ simulations within the respective bin) were highlighted to emphasise interaction configurations that poorly reproduced the metric of interest. A combination of these results (i.e., binned interaction configurations reproducing observed suppressiveness and observed alpha diversity) was then used to deduce the key interaction configuration(s) most likely to reproduce the observed characteristics of Mix-14 after antibiotics treatment (and *C. difficile* inoculation).

### Metabolic modelling (relevant for Fig. 3a)

Genome-scale metabolic models were reconstructed using CarveMe (v 1.5.1) [95] and gapfilled using KEMET (v 1.0.0) [96]. Community metabolic simulations were carried out using SMETANA (v 1.1.0) [97], with the --detailed and --molweight flags used to identify a minimal media and predict nutritional dependencies. 30 simulations were carried out in order to account for variability in the identified minimal media. Auxotrophies were calculated using the function “auxotrophies” in the module ‘cobra.auxotrophy’ of the ‘reframed’ Python package (v 1.4.0).

Alluvial diagrams were generated using the ‘ggalluvial’ R package (v 0.12.5). Heatmaps were generated using the ‘ComplexHeatmap’ R package (v 2.13.1) [98].

### Metabolomics data analysis (relevant for Fig. 4a)

The log-transformed, batch-normalized, and imputed untargeted metabolomics data was processed using R (v 4.1.0) and the ‘tidyverse’ package (v 2.0.0). The ‘Rstatix’ package (v 0.7.2) was used for statistical testing (Mann-Whitney U test), and the Benjamini-Hochberg procedure was used to control for false discovery rate due to multiple hypothesis testing.

### Mix-14 Polysaccharide utilization loci annotations (relevant for Fig. 4b)

The diamond sequence aligner (v 2.0.8, *diamond blastp*) was used to identify experimentally validated carbohydrate degrading hits from the dbCAN-PUL database [99] across the Mix-14 genomes in this study. Filtering criteria of sequence identity similarity of at least 60% and a bitscore of at least 100 were used to filter out spurious hits. The phylogenetic tree was reconstructed based on multiple sequence alignments of whole genome amino acid sequences using PhyloPhlAn (v3.0.67) [100]. The default PhyloPhlAn database was used with 32 cores for processing, alongside the --fast and --diversity low parameters.

### Mix-14 B3 *de novo* biosynthesis annotations (relevant for Fig. 4c)

Experimentally validated B3 biosynthesis genes were extracted from the database drafted by Magnusdottir and colleagues [101]. The database, obtained as a FASTA file, was curated using *seqtk* (r82) in order to remove corrupted entries; *cd-hit* (v 4.8.1) with default parameters (-c 0.9) was used to dereplicate the FASTA file [102]. Finally, the FASTA file was converted to a diamond database with *diamond makedb* and the genomes of interest were aligned on the database using *diamond blastp* (v 2.0.8) [103].

### Analysis of cohort metagenomes (relevant for Fig. 5a-c, f)

To assess the association of 14 core human gut bacteria in CDI conditions, an abundance mapping at the strain level was performed using publicly available gut metagenome sequencing datasets collected from fecal samples of individuals enrolled in clinical trials. For all the metagenome reads, quality trimming and adapter clipping were done with Trimmomatic [104]. Further, the reads were aligned against the human genome to filter out human reads and were assembled with Bowtie2 (v 2.3.2) [105] and SAMtools [106]. The resulting contigs were classified taxonomically by k-mer analysis using Kraken2 [107], with Kraken 2 standard database. Subsequent species abundance estimation was performed using Kraken-tools [108] with a threshold set to ignore species with fewer than 10 classified reads.

To assess the gene abundance for genes involved in FOS-generating, Stickland (precursor), B3 biosynthesis, and polyamine pathways, an abundance mapping was performed using metagenome sequencing datasets. Initially, quality trimming and adapter clipping were done with Trimmomatic. The trimmed reads were then assembled using metaSPAdes. The resulting contigs were subjected to sequence alignment using Diamond BLASTx against a custom database created from the FASTA sequences of the genes involved in the aforementioned pathways. The abundance data were compiled and filtered to include only hits with a bitscore greater than 100. Metadata were integrated, and the data were normalized and grouped by pathway to determine the gene abundance.

### Analysis of RBX2660 Mix Data (relevant for Fig. 5d-e)

Genus-level relative abundances of the RBX2660 Mix clinical trial [59] were filtered to exclude non-bacterial genera, after which the remaining genera for each patient were summed in their relative abundance 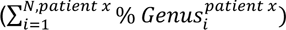, and then multiplied by 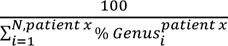 to correct relative abundance space after exclusion of non-bacterial genera (e.g., *Saccharomyces*). Then, patients for whom at least 50% of the relative abundance space was occupied by non-bacterial genera were excluded (i.e., 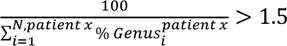) to discount unrepresentative/confounded bacterial compositions. In addition, we excluded patients also excluded by main authors as detailed in S1 [59]. The ratio was calculated between *Bacteroides* + 0.0001 and *Escherichia* + 0.0001, to avoid issues with log10-transformation of the ratio. Outlier ratios of >10^5 (log10 > 5) were excluded to discount artefacts of sequencing error at low to near-zero relative abundance space. Shannon alpha diversity was also calculated for each patient across bacterial relative abundances. To assess the role of either of these metrics in the post-treatment (Days > 0) data, statistical analyses were performed between treatment (placebo, single-, double-, open label) and outcome groups (i.e., binary variable of rCDI within 60 days) using Mann-Whitney U test. To account for false discovery rate in multiple comparisons, p-values were subsequently adjusted using the Benjamini-Hochberg (BH) procedure.

## DATA AVAILABILITY

Raw 16S sequencing data were uploaded onto the NCBI database under the Project ID code: PRJNA1136646. Metabolomics data were uploaded onto the Zenodo data repository under the accession code: DOI; 10.5281/zenodo.12968992

## CODE AVAILABILITY

Scripts used for statistical analysis and ecological modelling can be found in the following GitHub repository: https://github.com/NaomiIrisvdBerg/C_diff_project

## ACKNOWLEDGEMENTS

We thank Julia Nelson, Animal Disease Research and Diagnostic Laboratory, South Dakota for the help with 16S sequencing, and Metabolon, Inc. for the untargeted metabolomics. Metagenome data analysis for this project was performed using the OU Supercomputing Center for Education & Research (OSCER) at the University of Oklahoma (OU). We thank Suryang Kwak and Joohee Choi for their help in providing further details on their dataset: [59] This project has received funding from the United States National Institutes of Health (contract no: G-202306-70697), Walter R. Sitlington Endowment (grant no: 5-153200), European Research Council (ERC) under the European Union’s Horizon 2020 research and innovation programme (grant no. 866028) and from the UK Medical Research Council (project no. MC_UU_00025/11).

## AUTHOR CONTRIBUTIONS

AA and JS conceived the experimental study and planned all the *in vitro* experiments. AA, SM, AM and SG performed experimental work. NB conceived and carried out ecological modelling and contributed to all data analysis and figure composition. NB and KRP contributed to interpretation of modelling results and design of follow-up experiments and analyses. FZ and AB performed genome annotations and genome-scale metabolic modelling. NB, AA, and FZ contributed to metagenome annotation and analysis. LK helped with data analysis. FZ analyzed the metabolomics data. NB and AA performed the literature review and prepared the figures. NB, AA, JS and KRP wrote the manuscript. All authors read and/or commented on the manuscript.

## 4. SUPPLEMENTARY FIGURES

**Supplementary Figure 1.**
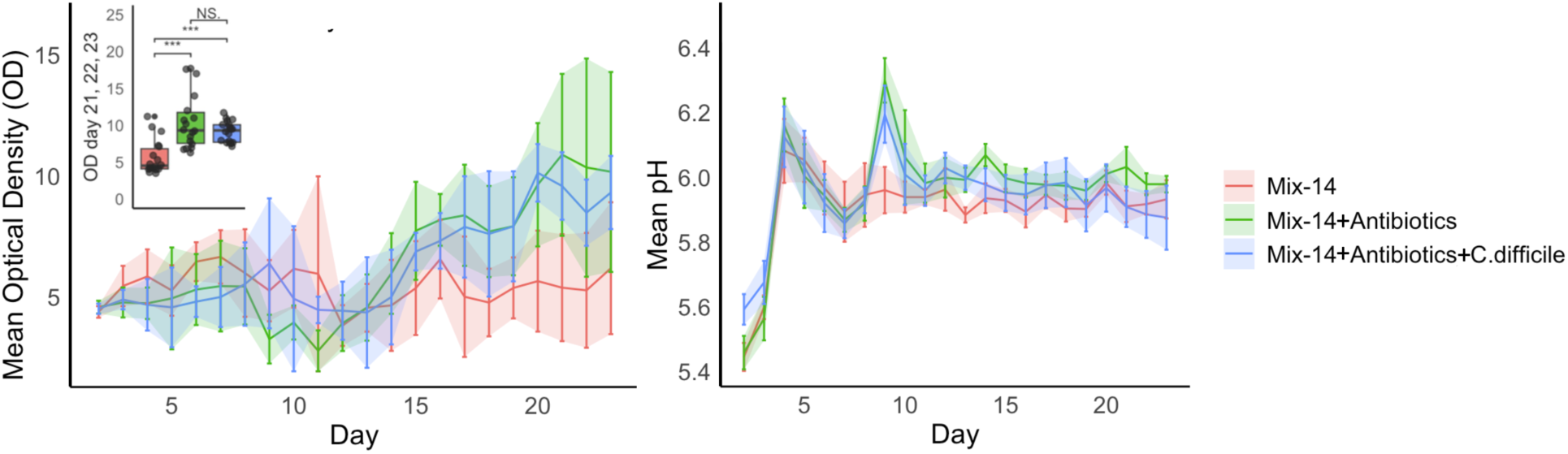
OD (left) and pH (right) trajectories of Mix-14 (control, in red), Mix-14 + Antibiotics (in green), and Mix-14 + Antibiotics + *C. difficile* (in blue). The inset of the left panel shows that final community OD values (for day 21, 22, 23) were significantly higher in antibiotics-treated Mix-14 than in Mix-14 control. Unlike OD, pH values showed no significant difference between treatment types. Error bars represent ± standard deviations from the mean.

**Supplementary Figure 2.**
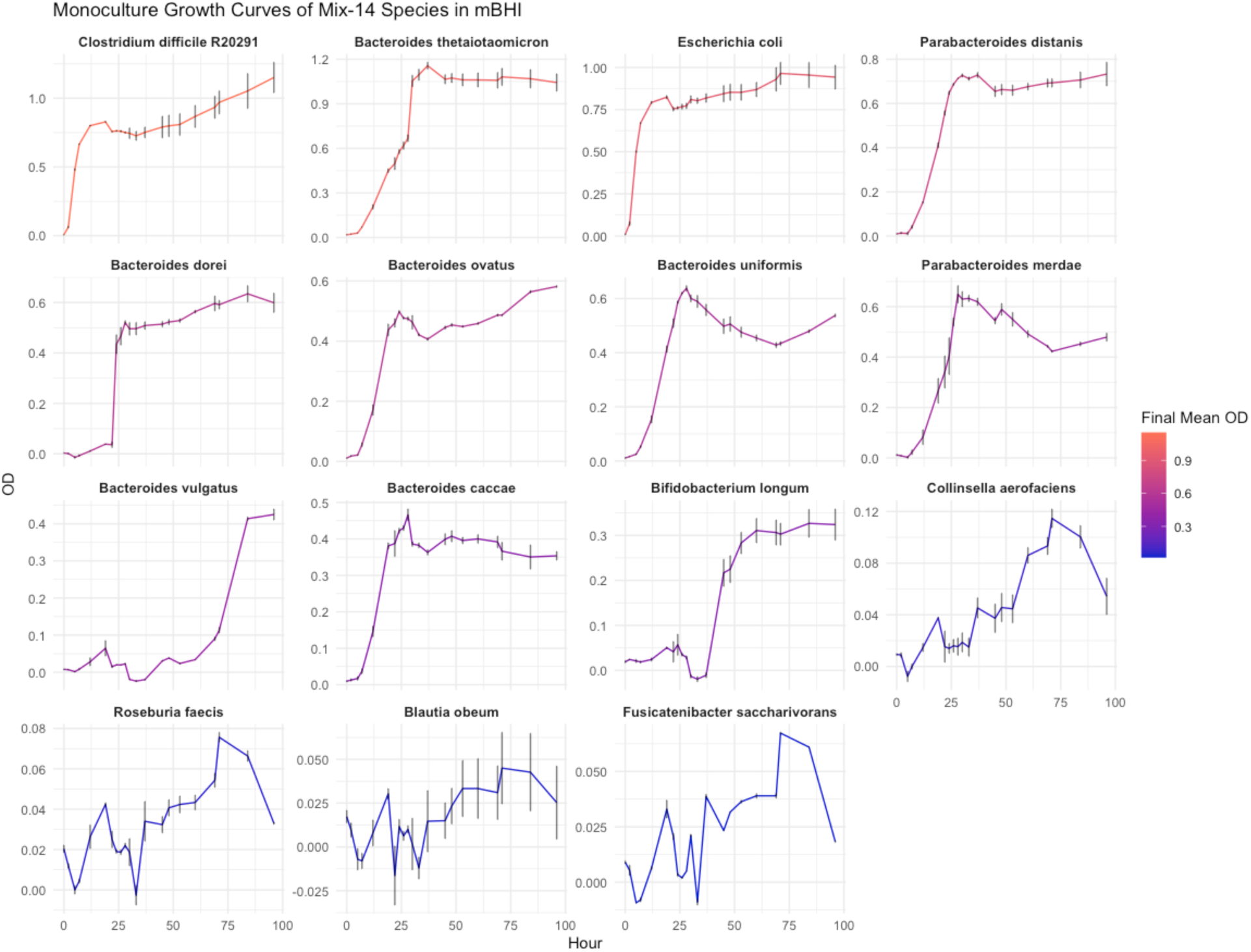
Monoculture growth curves for Mix-14 members. Note that *C. difficile* has the highest growth kinetics of all species, consistent with growth phenotypes of opportunistic pathogens. Error bars represent ± standard error from the mean (through which the line was fitted).

**Supplementary Figure 3.**
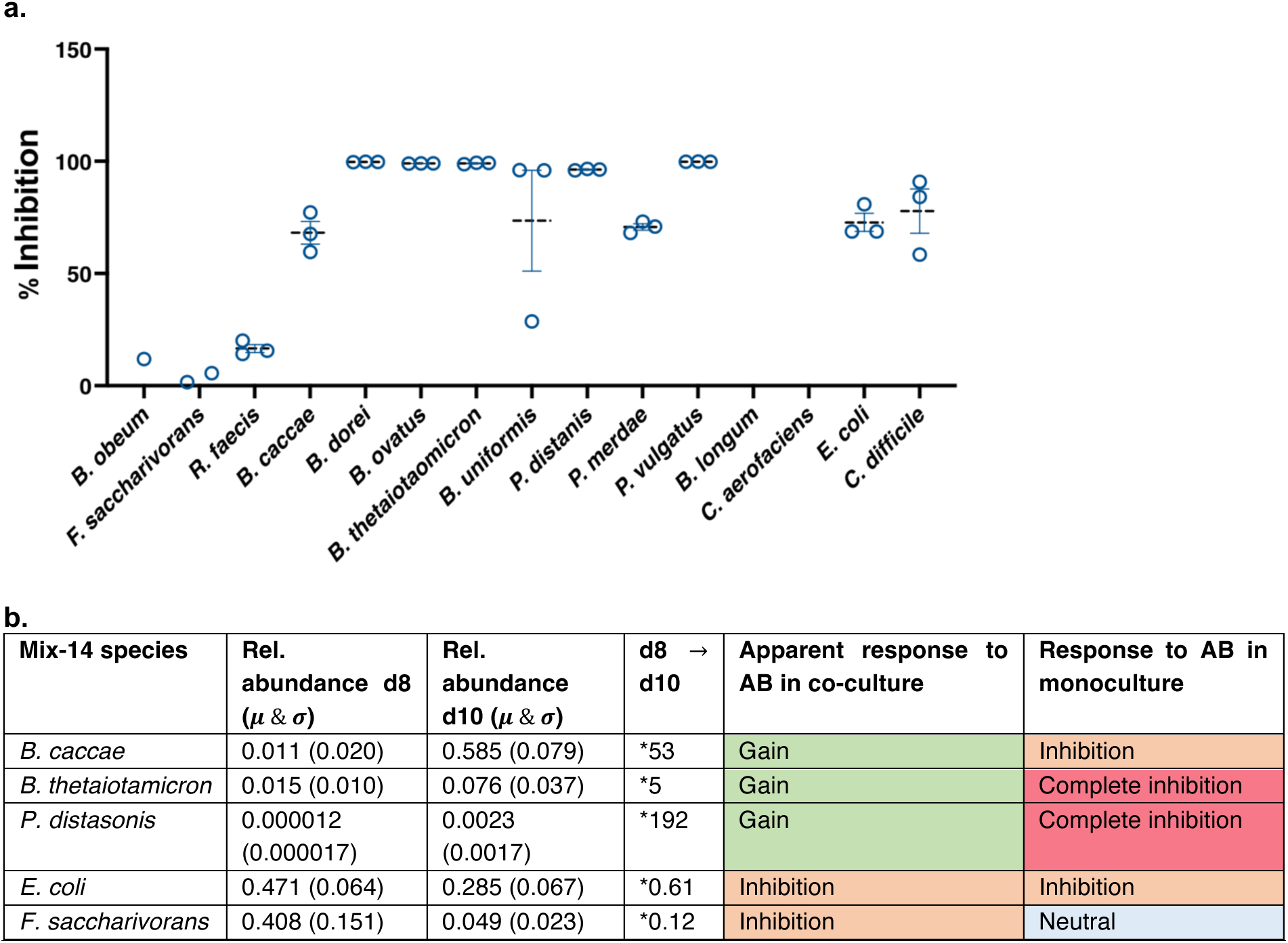
**a.** Individual antibiotics (clindamycin, at 250 𝜇g/mL concentration) sensitivity measured for Mix-14 members. **b.** Table contrasting antibiotics sensitivity of key Mix-14 species as measured in monoculture versus their apparent response in co-culture. For most key species, the response to the antibiotics in monoculture is unparalleled or even opposite to the apparent response in co-culture, emphasising community interaction effects such as cross-protection [109] and/or cross-sensitization [41].

**Supplementary Figure 4.**
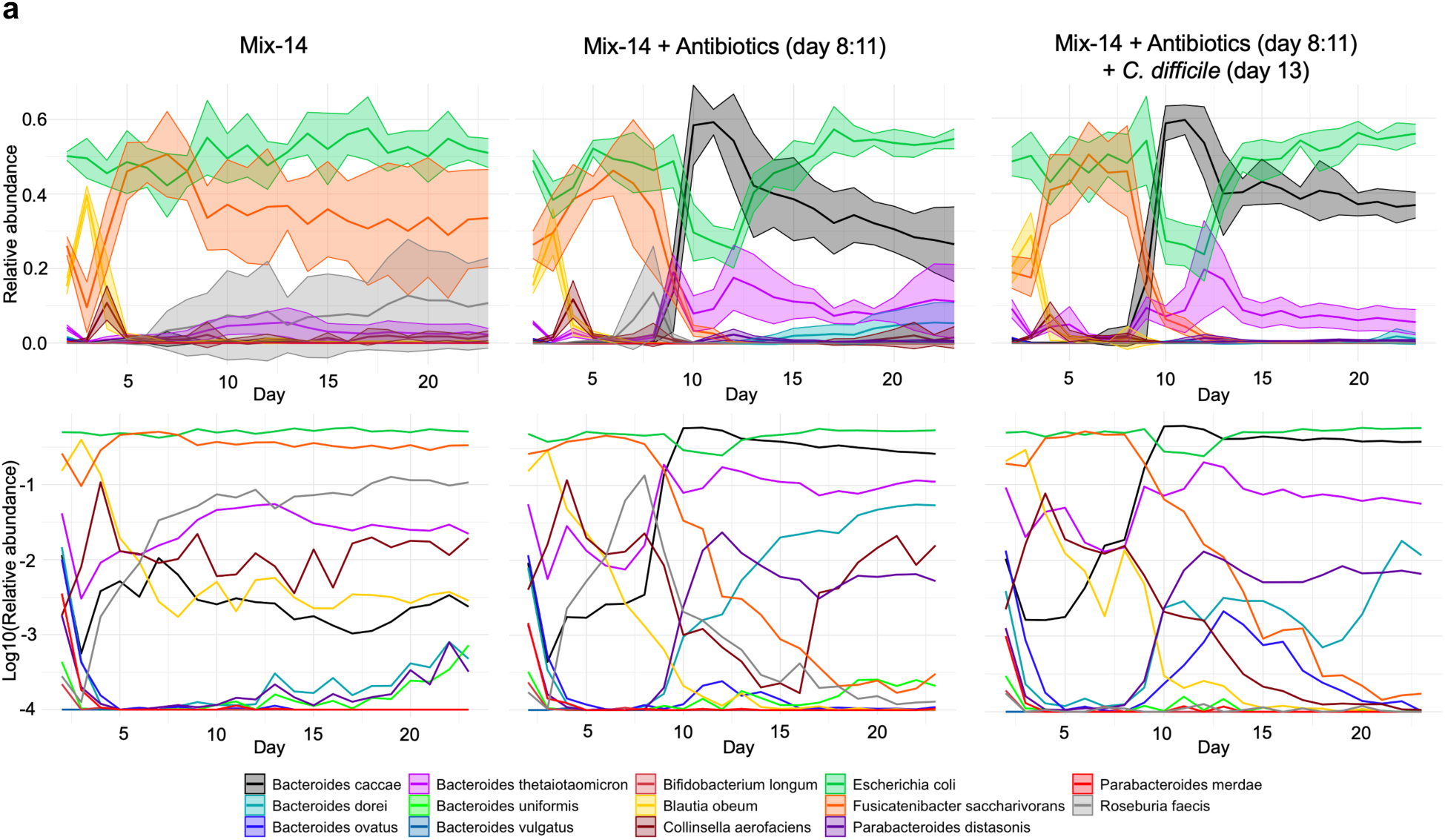
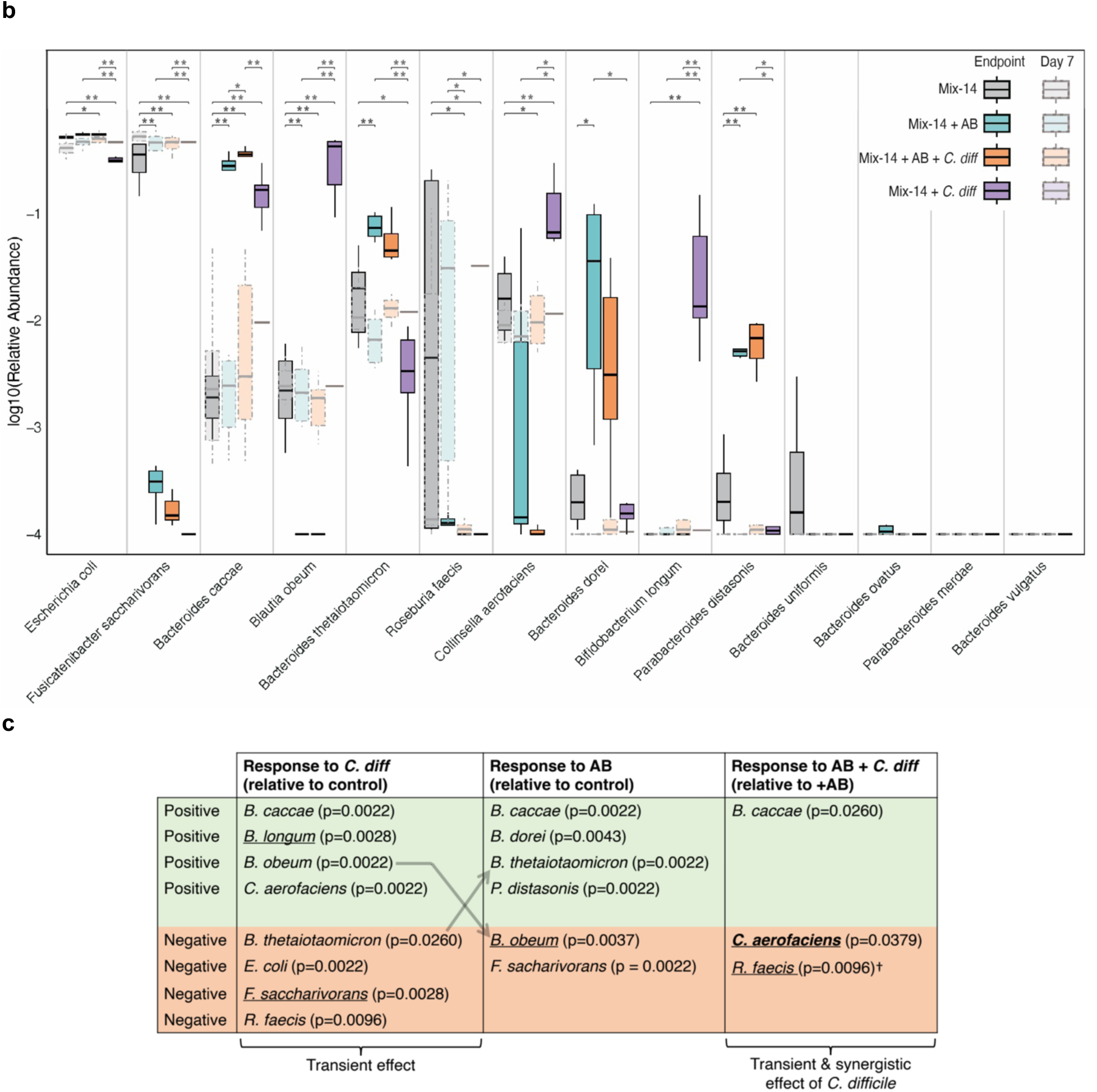
**a.** Relative abundance trajectories of Mix-14 species across different experimental conditions. Upper panels show mean relative abundance (solid lines) with ± standard deviation (shaded area), while bottom panels show log10-transformed mean relative abundances to emphasise trajectories of species in lower abundance regimes. **b.** Final and Day 7 species relative abundances compared between conditions, with Day 7 corresponding to the pre-perturbation timepoint (i.e., when growth conditions between different experimental conditions were identical, aiding to understand robustness of Mix-14 assembly). Statistical comparisons of endpoint data were done using Mann-Whitney U-test and subsequent BH-adjustments of p-values, and comparisons shown are those that were significant. Statistical significance is indicated as follows: *p < 0.05, **p < 0.01. Boxplot center lines represent respective medians, box limits represent upper and lower quartiles, and whiskers represent 1.5x interquartile range. **c.** Table summarising statistically significant responses of individual Mix-14 community members to the different perturbation regimes, as detailed in Fig. 2b. Statistical comparisons of species relative abundances were performed using the Wilcoxon rank-sum test (Mann-Whitney U test). Underlined = species (nearly) strictly dependent on or inhibited by perturbation. **Bold** = species’ response to combined disturbance is opposite to its response to either disturbance alone. Arrows highlight opposite responses by species to either perturbation. †= Effect could be due to load difference pre-perturbation (day 7). While strict sensitivities (see underlined) hint at direct effects of the perturbation, non-strict sensitivities may hint at indirect effects of the perturbation (i.e., via effects on the interaction landscape, which are of importance in Mix-14 as emphasised by the discrepancy between monoculture AB sensitivities and community-level sensitivities).

**Supplementary Figure 5.**
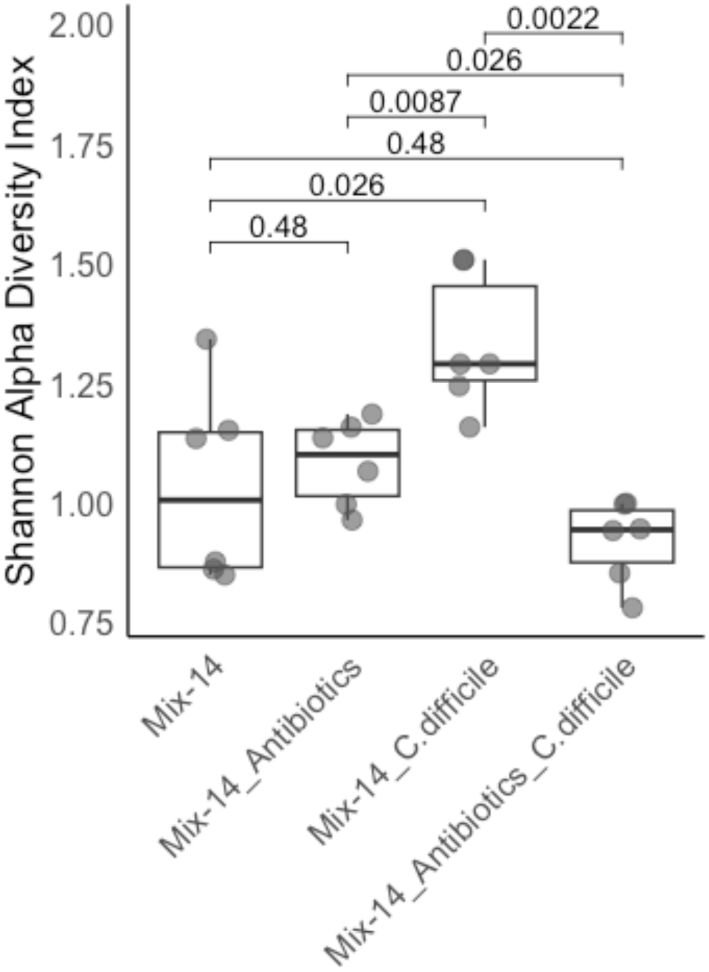
Comparison of Mix-14 Final Shannon alpha diversity across experimental conditions. Statistical comparisons (Mann-Whitney U, BH-adjusted) show that in the absence of antibiotics, *C. difficile* challenge induced community shifts that increased the system’s alpha diversity, despite the pathogen being suppressed by Mix-14, hinting at a community-wide transient effect. However, in combination with antibiotics, the *C. difficile* challenge significantly lowered alpha diversity relative to antibiotics alone. Boxplot center lines represent respective medians, box limits represent upper and lower quartiles, and whiskers represent 1.5x interquartile range

**Supplementary Figure 6.**
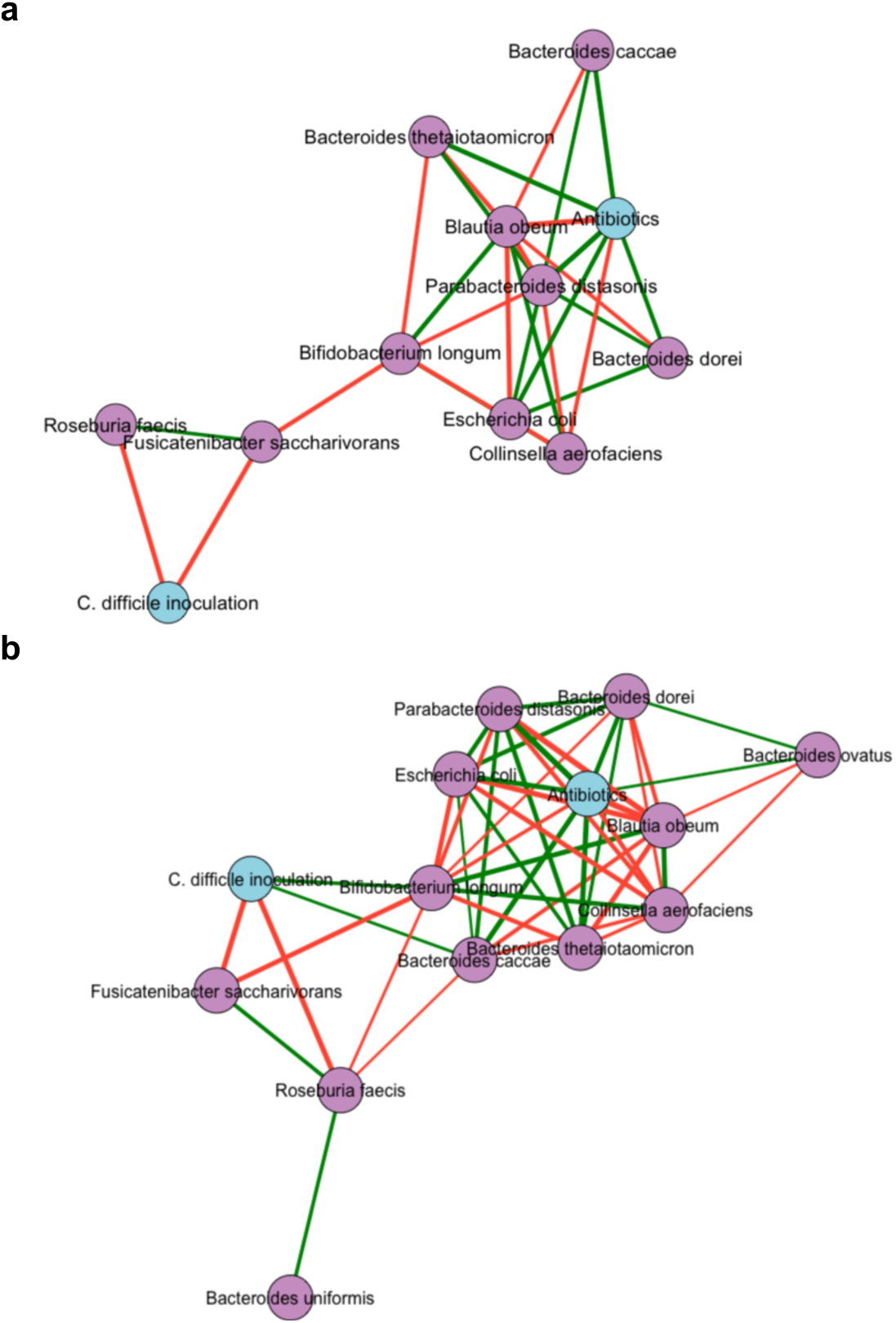
Correlation networks of end-point relative abundances of all four conditions, with presence (1)/absence (0) values assigned to *C. difficile* and antibiotics treatment (nodes in blue). Mix-14 (control) thus has 0 assigned to either disturbance variable, while Mix-14 + Antibiotics + *C. difficile* has 1 assigned to both. Correlations emphasize potential directions of net interaction effects. **a.** Spearman correlations, p<0.05, rho >|0.6|. **b.** Spearman correlations, p<0.05, rho >|0.4|.

**Supplementary Figure 7.**
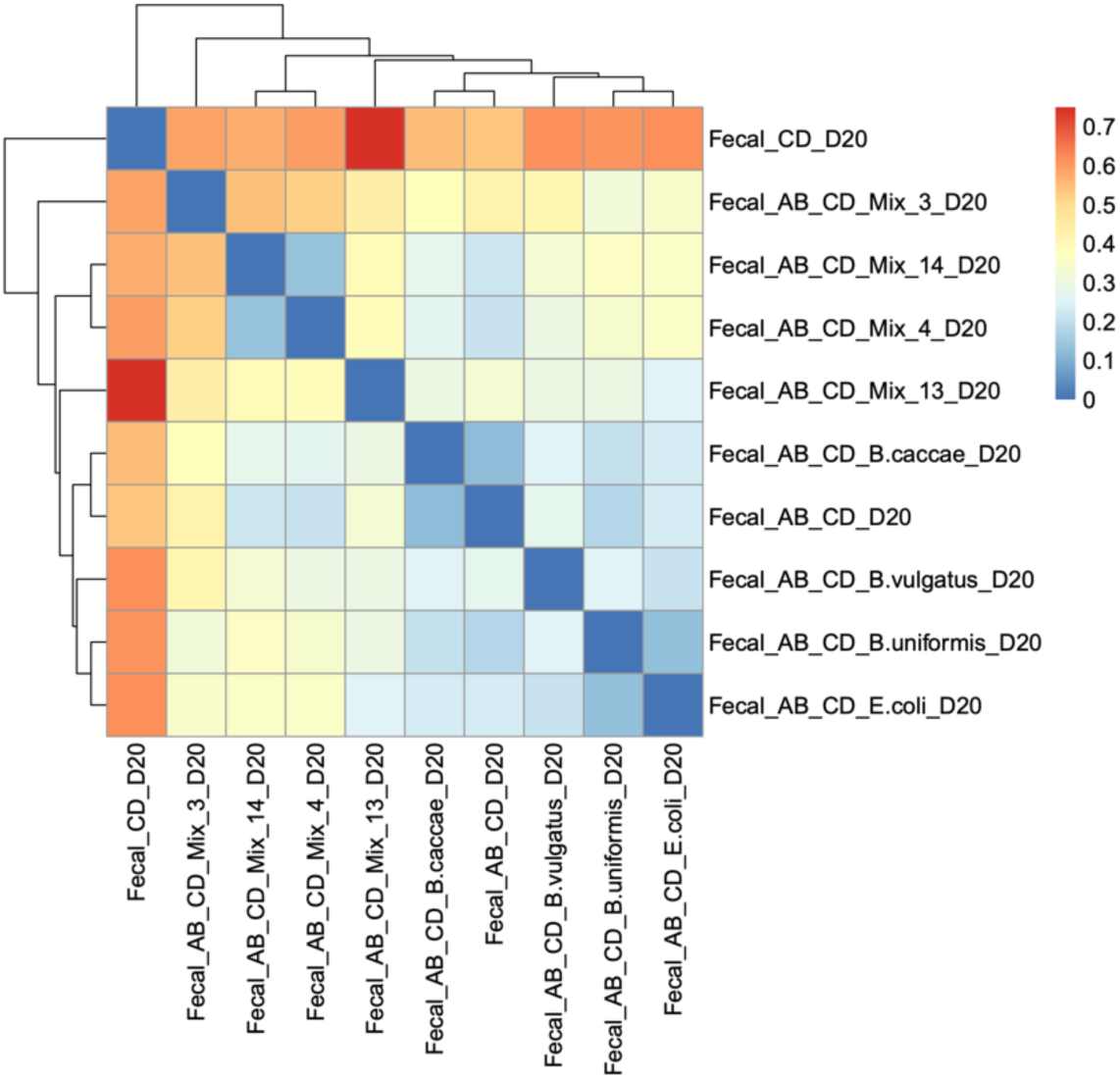
Bray-Curtis dissimilarities of endpoint (Day 20) compositions of fecal-matter derived communities. Most dissimilar in this comparison is ‘Fecal_CD_D20’ – the only community not treated with antibiotics. Suppressive treatments (e.g., ‘Fecal_AB_CD_Mix_14_D20’, ‘Fecal_AB_CD_Mix_4_D20’) and the ‘Fecal_CD_D20’ control tend to cluster together, suggesting that similar community compositions might be crucial for suppression of *C. difficile*. Conversely, non-suppressive treatments tend to be clustered away from suppressive ones, reinforcing the notion that their community compositions diverge in ways that might affect their functional outcomes against *C. difficile*.

**Supplementary Figure 8.**
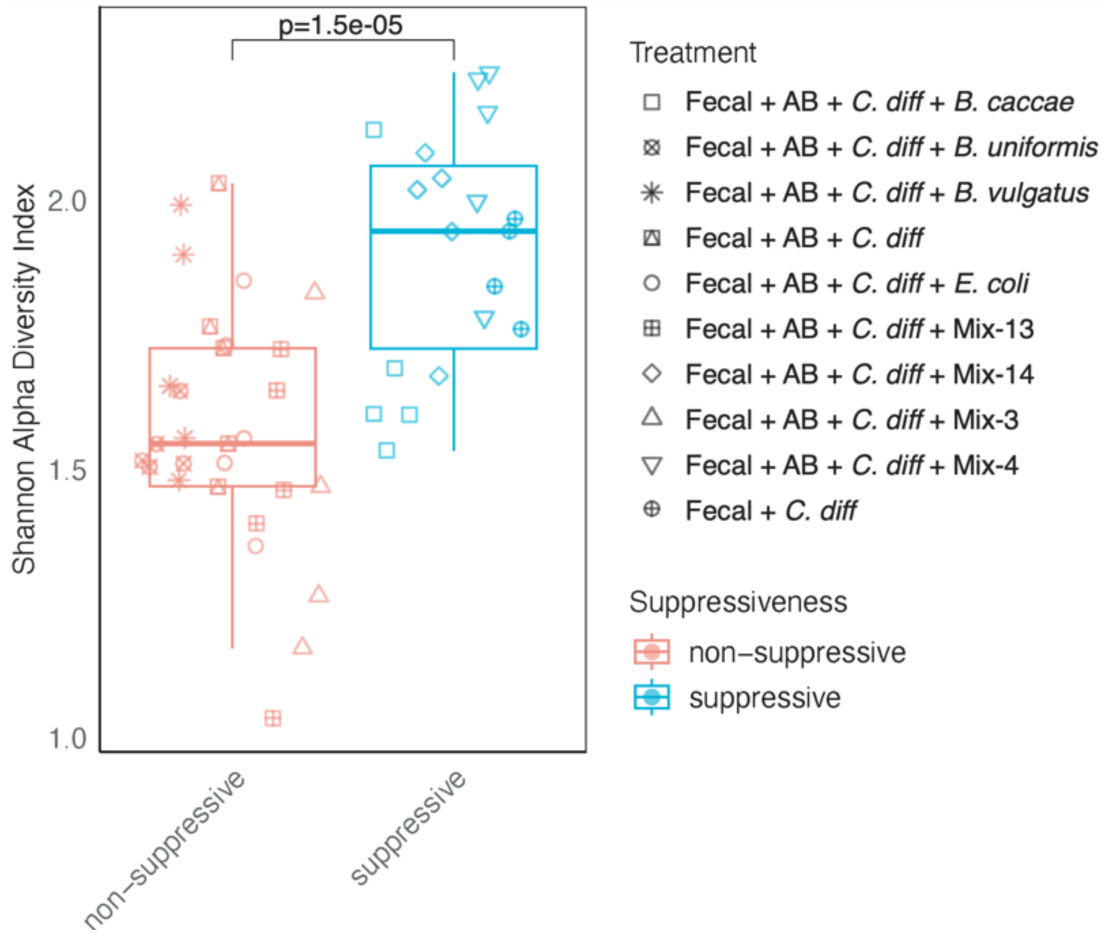
Comparison of Shannon alpha diversities of suppressive fecal-matter derived communities versus non-suppressive ones. Suppressive fecal-matter derived communities have significantly higher (Mann-Whitney U, p<0.05) alpha diversity than non-suppressive communities. Boxplot center lines represent respective medians, box limits represent upper and lower quartiles, and whiskers represent 1.5x interquartile range.

**Supplementary Figure 9.**
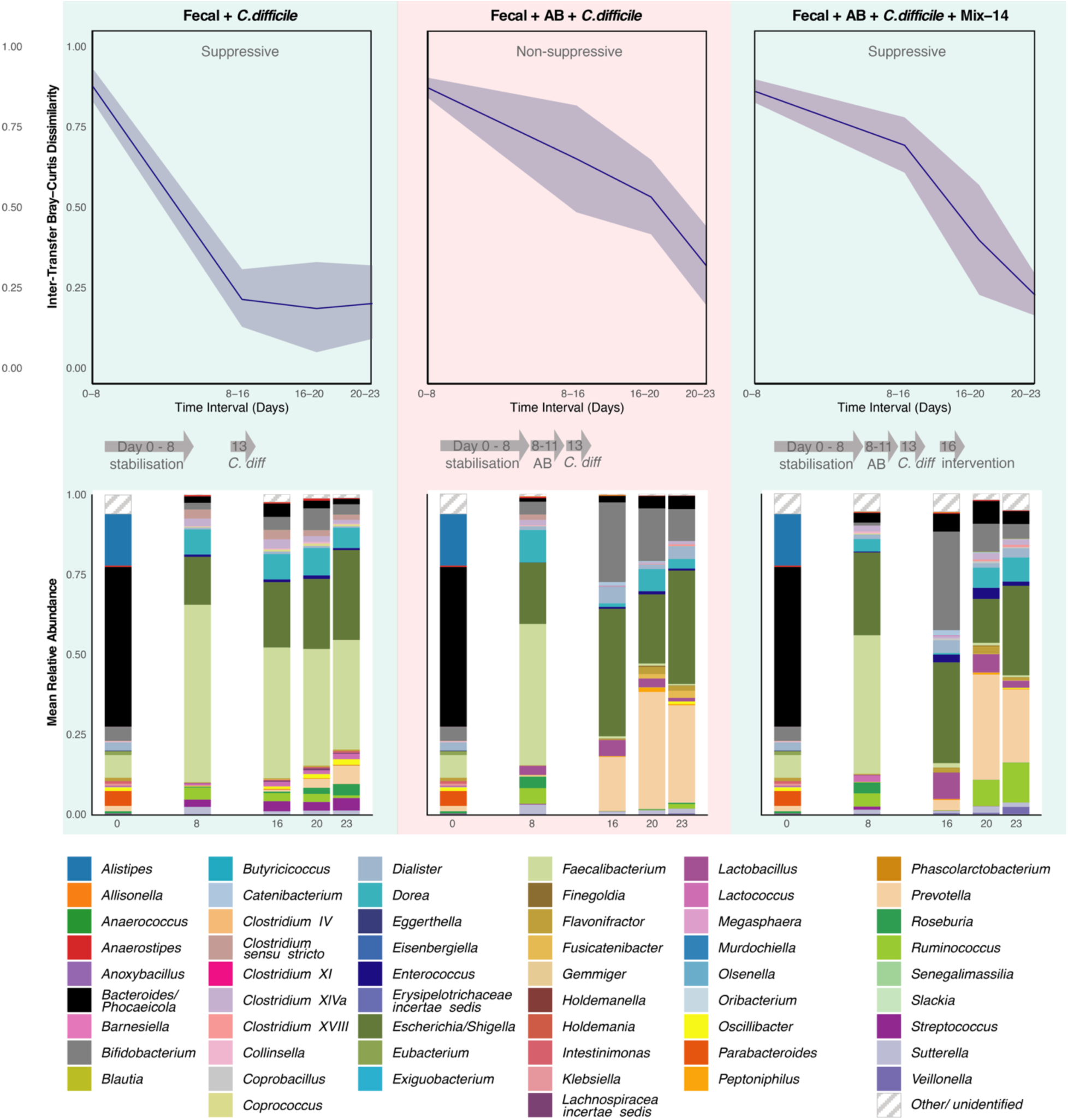
Inter-transfer Bray-Curtis dissimilarity (top half, mean & ± standard deviations) and genus-level relative abundance distribution (bottom half) of fecal-matter derived communities challenged with *C. difficile* (left panels), treated with antibiotics (AB) and subsequently challenged with *C. difficile* (middle panels), or perturbed by both antibiotics, subsequently challenge with *C. difficile,* and then treated with a Mix-14 intervention (right panels). As in line with Suppl. fig. 7, antibiotics induce significant shifts within the community, resulting in high inter-transfer Bray-Curtis dissimilarity, while the fecal community unperturbed by antibiotics shows rapid stabilisation (Day 0-8). Also noteworthy is the difference in composition of the fecal inoculum (see stacked barplot at Day 0) compared to the timepoints taken after growth in mBHI, corresponding to the expected effect of growing a bacterial community in a synthetic setting versus the setting of origin (the fecal environment).

**Supplementary Figure 10.**
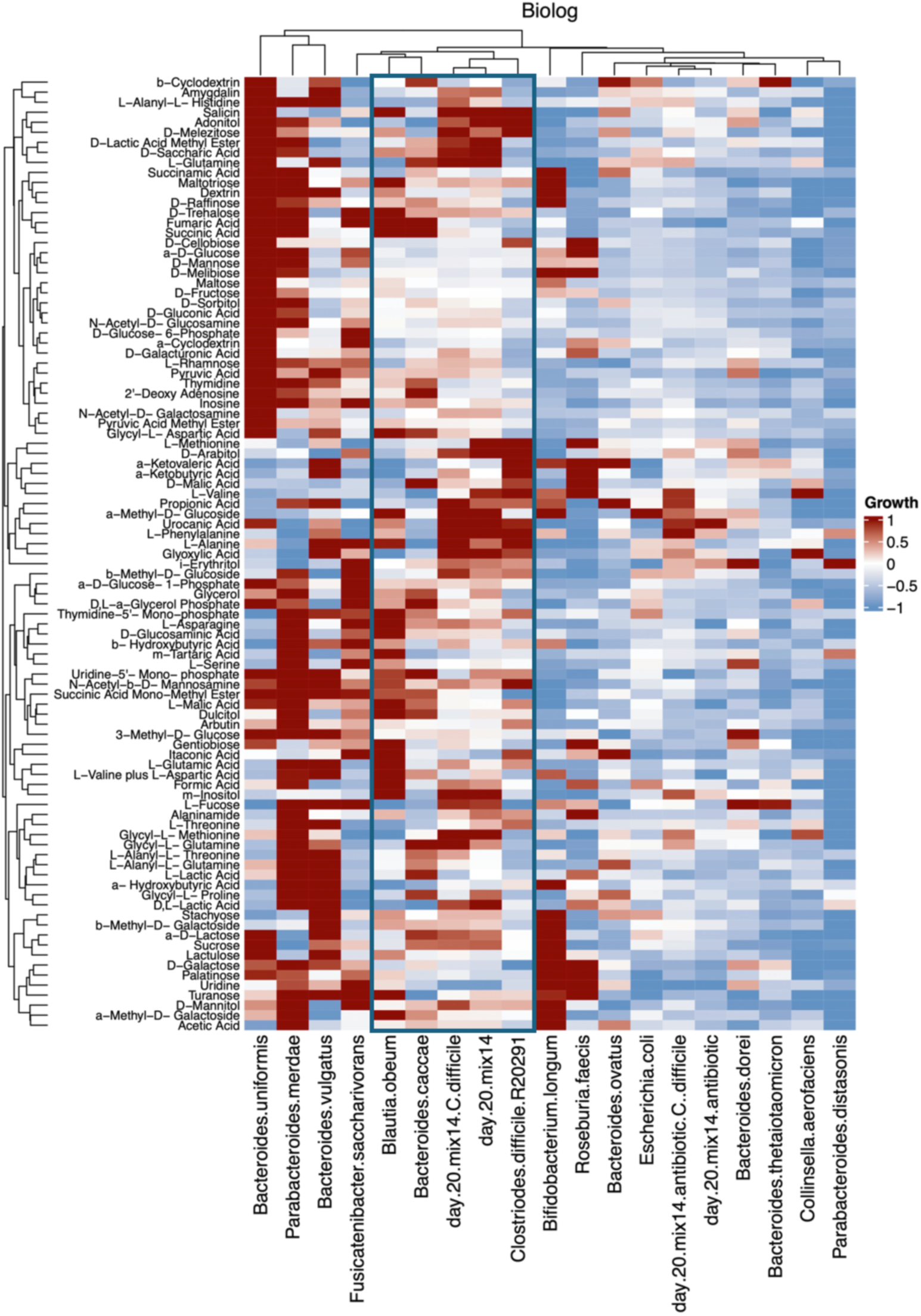
Pearson correlation heatmap comparing carbon usage assay profile similarities of Mix-14 species and different experimental conditions. The redder the colour, the more stimulated the growth (normalized against growth without the compound in question). Of particular interest in the Biolog data heatmap/cladogram is the highlighted branch containing *B. caccae* and *C. difficile*, containing other (individual) strains with similar carbon usage profiles - which could hence compete with *B. caccae* over shared carbon sources. In this similarity clade, only *B. caccae* does well in community with antibiotics treatment. This could indicate that *B. caccae* has a competitive upper hand over *C. difficile* and its approx. equidistant branch member *F. saccharivorans* for the shared resources following antibiotics (or *C.* difficile) perturbation.

**Supplementary Figure 11.**
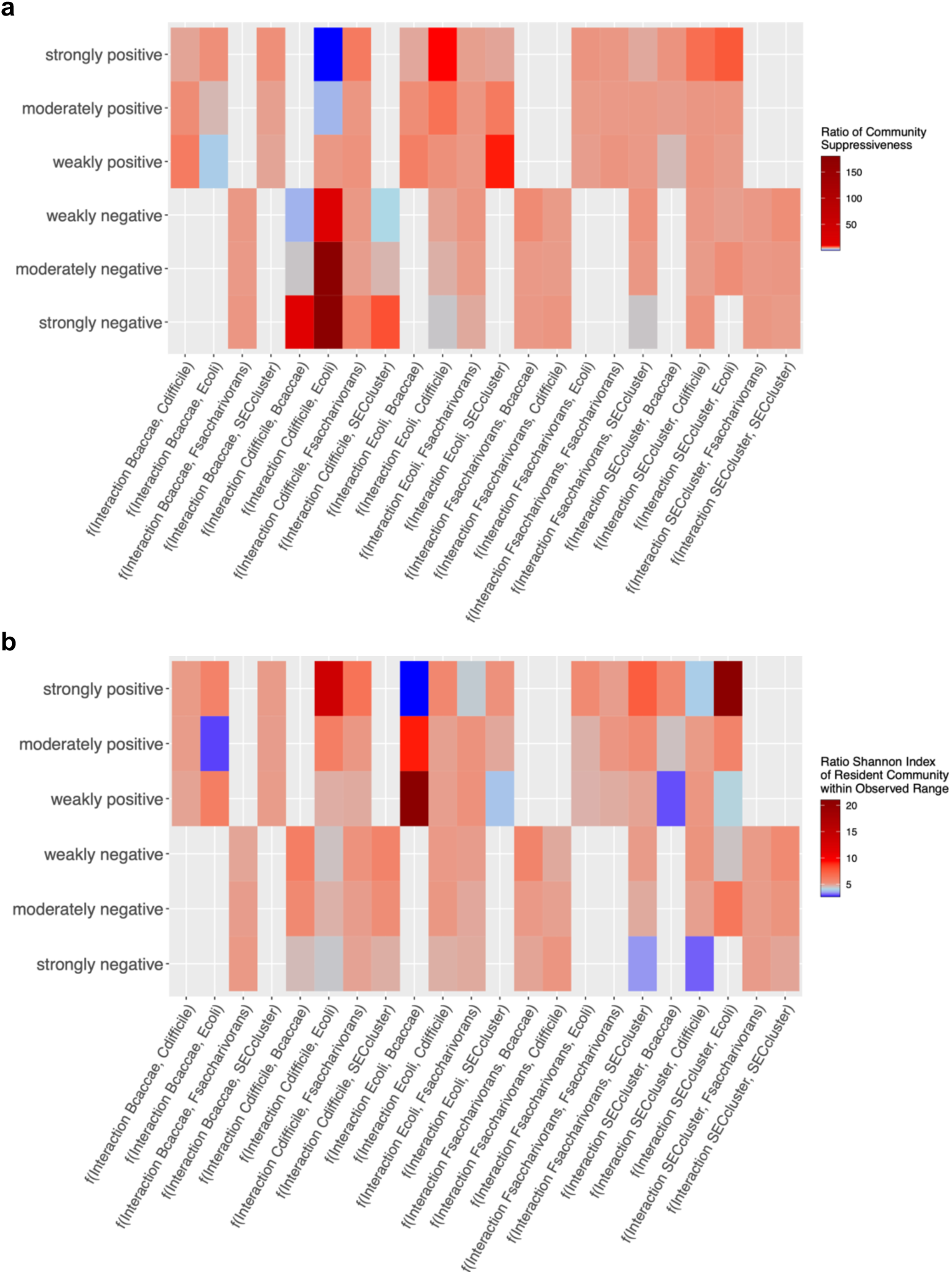
Ecological model simulation results. **a.** Ratio of suppressive versus nonsuppressive simulated final community states per binned interaction strength, across all binary interaction coefficients. A simulation was considered suppressive if the final abundance of *C. difficile* was less than its initial load. **b.** Ratio of Shannon diversity of the simulated final community states being within the observed range per binned interaction strength, across all binary interaction coefficients. For either heatmap, the higher the ratio, the more accurate the simulation results were for each specific binned interaction strength in reproducing the community characteristic (Suppressiveness, Shannon alpha diversity) observed experimentally for Mix-14 after antibiotic treatment. For the Suppressiveness scores, the colour gradient (blue, lightblue, red, darkred) used was set at 0, 1, 5, 10, whereas for the Shannon index range score, the colour gradient used was set at 0, 1, 3, 10 (due to a lower maximum ratio for the latter).

**Supplementary Figure 12.**
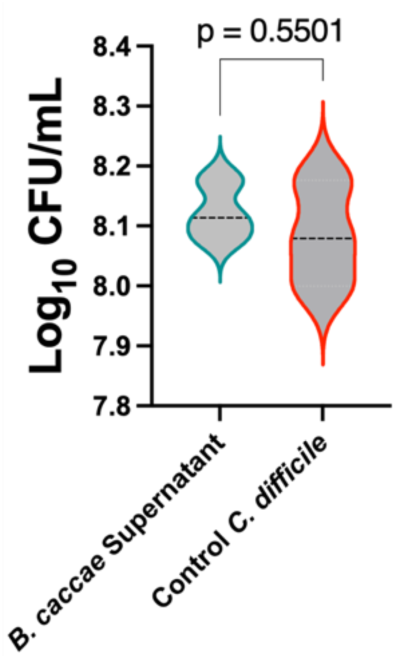
While *B. caccae* was shown to be critical in the *C. difficile* suppression conferred by Mix-14, its own direct suppressiveness (see Fig. 1b; *B. caccae* suppressed *C. difficile* by a little over one order of magnitude) – while significant – was not nearly as strong as that of the consortium (Fig. 2a, Mix-14, especially post antibiotics, suppressed *C. difficile* by over 6 orders of magnitude). Here, it is shown that *B. caccae*’s suppression is likely not mediated by antimicrobials or other antagonistic extracellular compounds it secretes, since *C. difficile* load in its supernatant was not significantly lower than *C. difficile* load of control. Statistical significance was calculated using a two-tailed unpaired t-test.

**Supplementary Figure 13.**
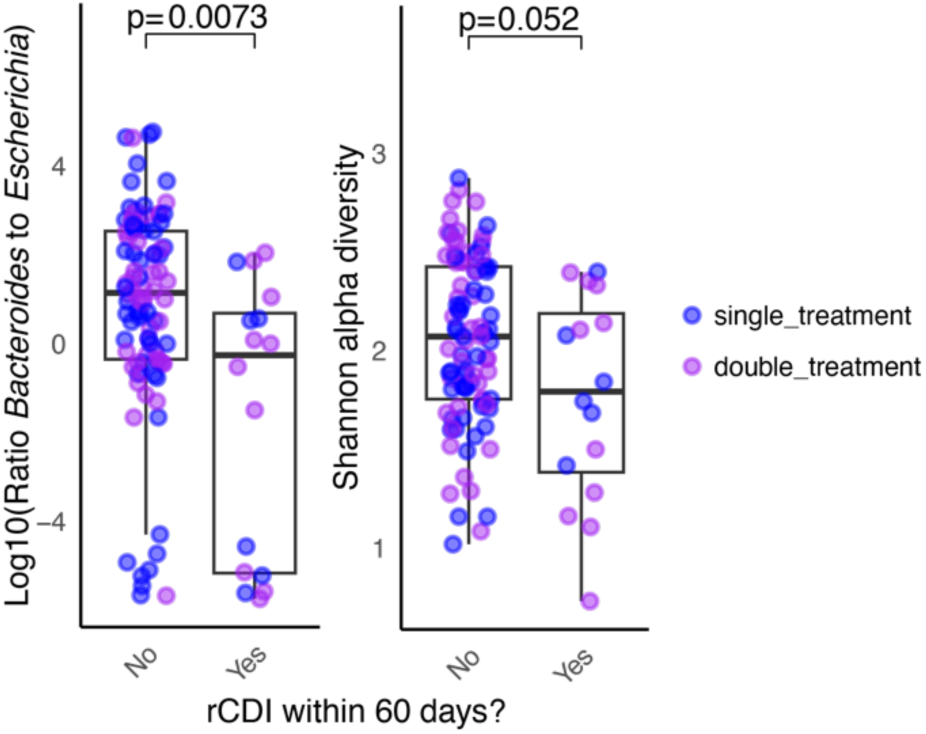
In addition to figure 5 panel e-d, we here show the comparisons of communitywide metrics when patients treated with single and double doses of probiotic mix RBX2660 [59] were grouped together. Log10 ratio of *Bacteroides* to *Escherichia* is significantly lower in failed treatments (t-test, p = 0.0073), while alpha diversity does not distinguish with the same statistical significance or power (t-test, p = 0.052). Median Ratio for successful treatments was 14.0 (1.15 in log10), while median Ratio for failed treatments was 0.651 (−0.26 in log10). Median Shannon alpha diversity for successful treatments was 2.07, and 1.79 for unsuccessful treatments. Boxplot center lines represent respective medians, box limits represent upper and lower quartiles, and whiskers represent 1.5x interquartile range.

**Supplementary Figure 14.**
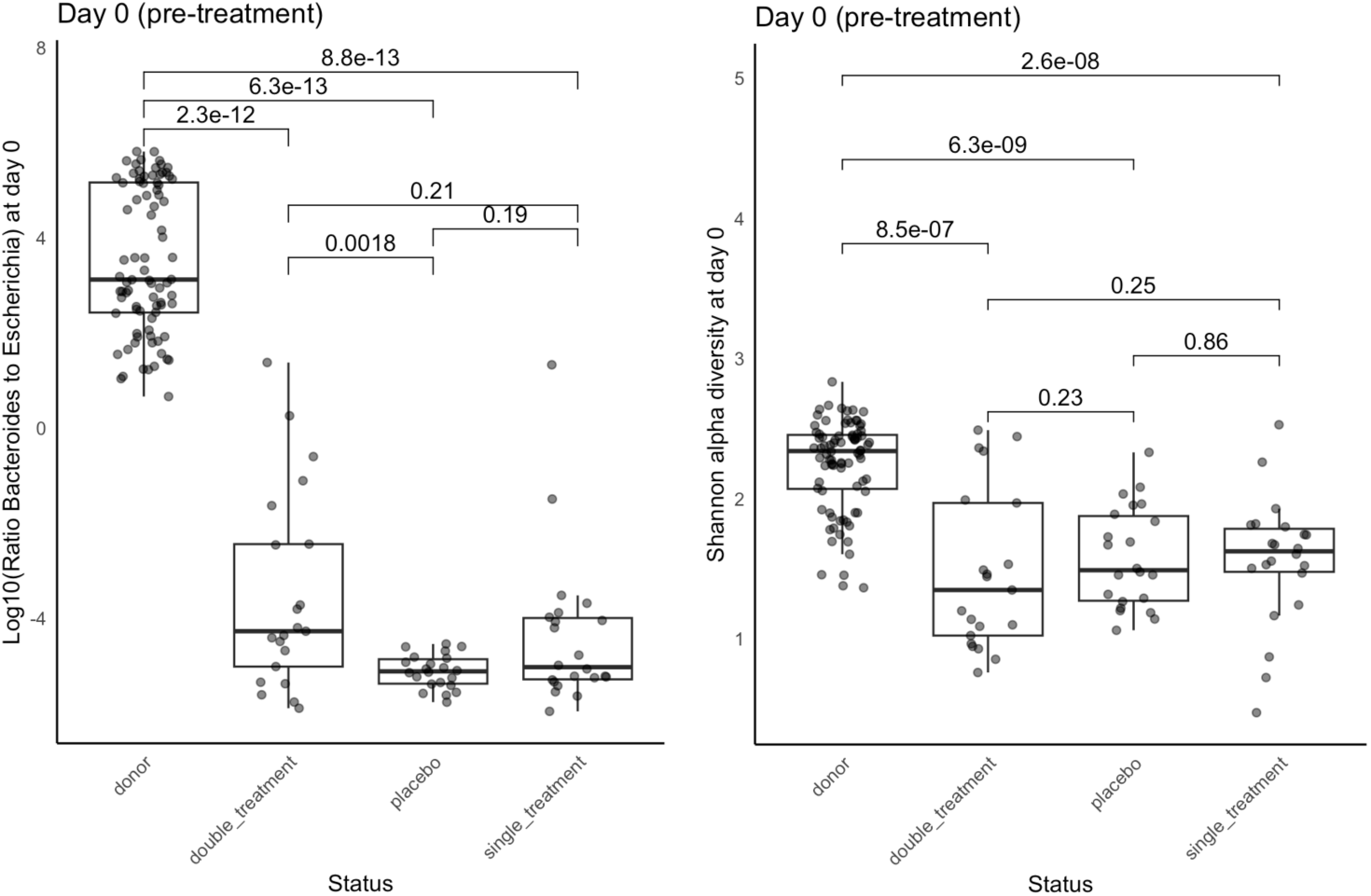
Donors from which strains were isolated and then prepared to formulate the RBX2660 mix for standardized live bacterial suspension [59], are defined by a significantly (Mann-Whiteny U test, BH-adjusted p-value < 0.05) higher ratio of *Bacteroides* to *Escherichia* (left panel) and Shannon alpha diversity calculated over Genus-level (right panel) than all the CDI patients that were divided among ‘placebo’, ’single treatment’ and ‘double treatment’ groups. At day 0, there were no differences in conditions for single treatment, double treatment, and placebo groups. While we do not see any significant differences in alpha diversity between these groups, we do see a significant difference in ratio between double treatment and placebo, with a subset of ‘double’ treatment group datapoints having a slightly higher ratio. Boxplot center lines represent respective medians, box limits represent upper and lower quartiles, and whiskers represent 1.5x interquartile range.

**Supplementary Figure 15.**
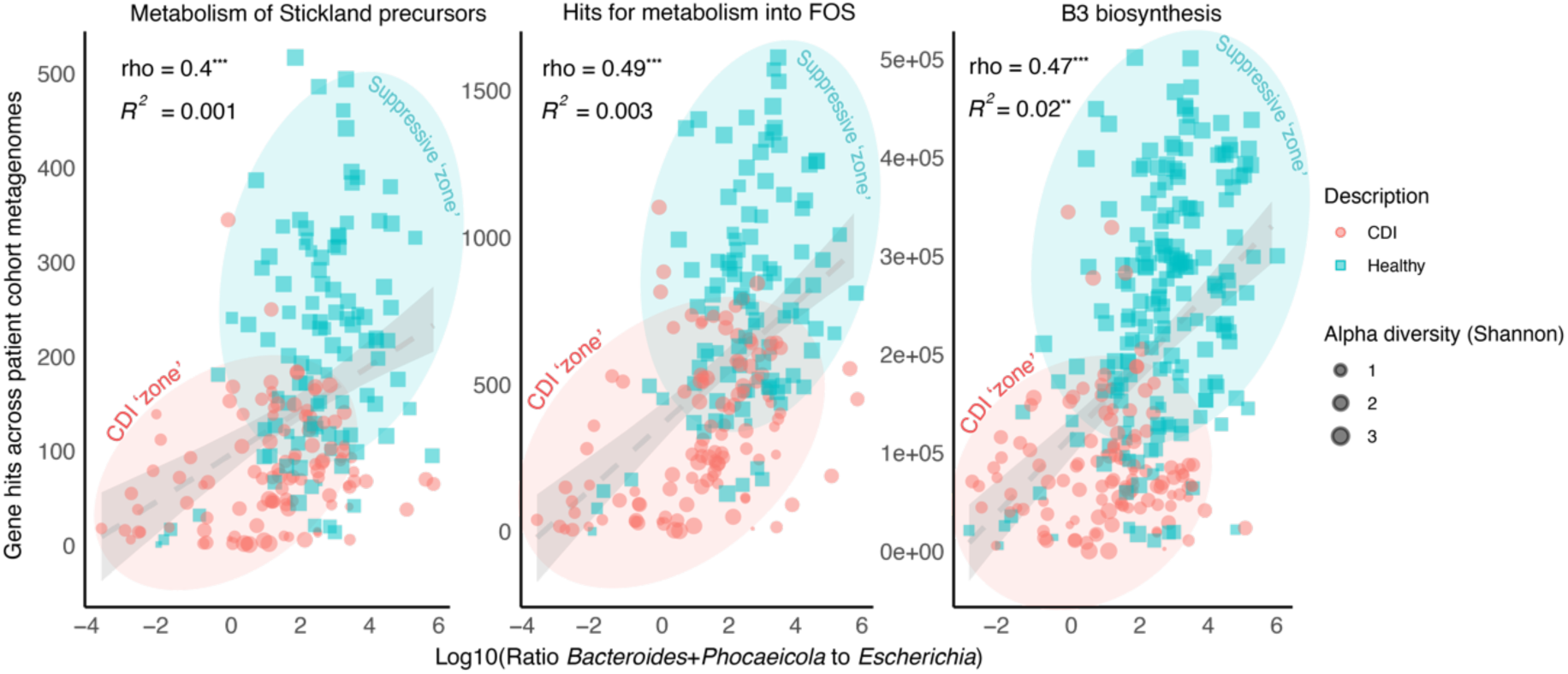
The ratio of *Bacteroides* to *Escherichia* and hypothesised molecular signatures – including B3 biosynthesis – are positively correlated, as was predicted from our *in vitro* data (Fig. 4). Here, we plotted the gene hits across metagenomic data of healthy individuals and CDI patients (excluding FMT patients). The fitted line represents the linear regression, and the shaded area the associated standard error.

## Supplementary Tables

**Supplementary Table 1:**
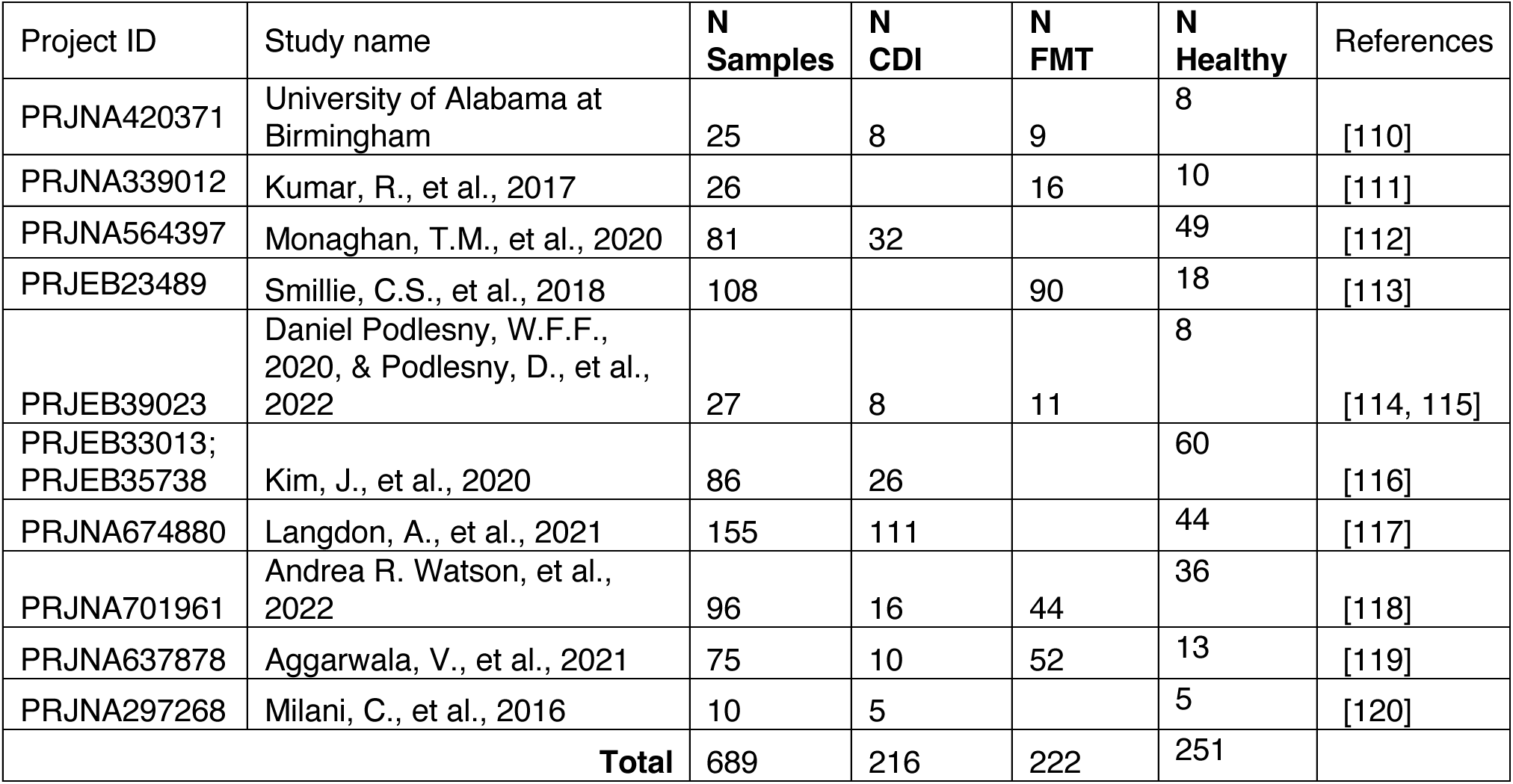
List of cohort studies included for metagenomic analysis. Sample numbers shown are those that met sequence quality criteria as mentioned in Methods (section 4.2.8)

**Supplementary Table 2:** Correlations of significant ratios with Shannon-Indexed Alpha Diversity. Combinations in grey have worse regression than the focus Ratio of *Bacteroides* to *Escherichia.* In bold is the ratio with the strongest regression.

| Ratio | Linear regression metrics |  | Spearman correlation metrics |  |
| --- | --- | --- | --- | --- |
|  | R<sup>2</sup> | p-value | rho | p-value |
| *Bacteroides*_Abundance / *Escherichia*_Abundance* | 0.437 | 3.152e-07 | 0.6892097 | 2.041e-07 |
| *Bacteroides*_Abundance / *Fusicatenibacter*_Abundance* | 0.3401 | 1.365e-05 | 0.6013895 | 9.576e-06 |
| (*Fusicatenibacter*_Abundance + *Parabacteroides*_Abundance) / *Bacteroides*_Abundance | 0.4198 | 6.424e-07 | -0.6448111 | 1.491e-06 |
| (*Escherichia*_Abundance + *Roseburia*_Abundance) / *Bacteroides*_Abundance | 0.4399 | 2.792e-07 | -0.6892097 | 2.041e-07 |
| (*Escherichia*_Abundance + *Fusicatenibacter*_Abundance) / *Bacteroides*_Abundance | 0.4409 | 2.675e-07 | -0.6873643 | 2.217e-07 |
| (*Collinsella*_Abundance + *Parabacteroides*_Abundance) / *Bacteroides*_Abundance | 0.4843 | 3.987e-08 | -0.6561007 | 9.01e-07 |
| (*Collinsella*_Abundance + *Fusicatenibacter*_Abundance) / *Bacteroides*_Abundance | 0.3416 | 1.289e-05 | -0.5687147 | 3.437e-05 |
| (*Collinsella*_Abundance + *Escherichia*_Abundance) / *Bacteroides*_Abundance | 0.4405 | 2.717e-07 | -0.6892097 | 2.041e-07 |
| (*Blautia*_Abundance + *Fusicatenibacter*_Abundance) / *Bacteroides*_Abundance | 0.3752 | 3.705e-06 | -0.6021494 | 9.283e-06 |
| (*Blautia*_Abundance + *Escherichia*_Abundance) / *Bacteroides*_Abundance | 0.4454 | 2.211e-07 | -0.6892097 | 2.041e-07 |
| (*Bifidobacterium*_Abundance + *Fusicatenibacter*_Abundance) / *Bacteroides*_Abundance | 0.589 | 1.976e-10 | -0.7625923 | < 2.2e-16 |
| **(*Bifidobacterium*_Abundance + *Escherichia*_Abundance) / *Bacteroides*_Abundance** | **0.632** | **1.502e-11** | **-0.7859314** | **< 2.2e-16** |
| (*Bifidobacterium*_Abundance + *Collinsella*_Abundance) / *Bacteroides*_Abundance | 0.5948 | 1.416e-10 | -0.7667173 | < 2.2e-16 |
| (*Bifidobacterium*_Abundance + *Fusicatenibacter*_Abundance) / *Blautia*_Abundance | 0.3692 | 4.659e-06 | -0.6060573 | 7.905e-06 |
| (*Bifidobacterium*_Abundance + *Fusicatenibacter*_Abundance) / *Collinsella*_Abundance | 0.2755 | 0.0001283 | -0.663591 | 6.439e-07 |
| (*Bifidobacterium*_Abundance + *Fusicatenibacter*_Abundance) / *Parabacteroides*_Abundance | 0.4142 | 8.048e-07 | -0.6574034 | 8.499e-07 |
| *(Bacteroides_Abundance + Roseburia_Abundance) / Escherichia_Abundance* | 0.4135 | 8.293e-07 | 0.6643508 | 6.223e-07 |
| *(Bacteroides_Abundance + Parabacteroides_Abundance) / Escherichia_Abundance* | 0.4194 | 6.527e-07 | 0.6861702 | 2.339e-07 |
| *(Bacteroides_Abundance + Roseburia_Abundance) / Fusicatenibacter_Abundance* | 0.3039 | 4.899e-05 | 0.5618758 | 4.428e-05 |
| *(Bacteroides_Abundance + Parabacteroides_Abundance) / Fusicatenibacter_Abundance* | 0.3161 | 3.205e-05 | 0.5755536 | 2.655e-05 |
| *(Bacteroides_Abundance + Collinsella_Abundance) / Escherichia_Abundance* | 0.425 | 5.186e-07 | 0.6764003 | 3.624e-07 |
| *(Bacteroides_Abundance + Blautia_Abundance) / Escherichia_Abundance* | 0.4218 | 5.909e-07 | 0.6761832 | 3.659e-07 |
| *(Bacteroides_Abundance + Collinsella_Abundance) / Fusicatenibacter_Abundance* | 0.3222 | 2.579e-05 | 0.5785931 | 2.364e-05 |
| *(Bacteroides_Abundance + Blautia_Abundance) / Fusicatenibacter_Abundance* | 0.3178 | 3.017e-05 | 0.5787017 | 2.354e-05 |

\* = inverse ratio has identical metrics, with inverse sign for correlation coefficient.

**Supplementary Table 3A:** Overview of parameters used in the ecological model (see Methods 4.2.2.)

| Parameter | Units | Description | Informed by |
| --- | --- | --- | --- |
| $r_i$ | $[d]^{-1}$ | Maximum per capita growth rate of bug $i$. | Max growth rate of monoculture growth data (mBhi_GC.xlsx), pulled from a normal distribution: $r_i \sim N(\mu(r_i), \sigma(r_i)^2)$ |
| $\phi$ | $[d]^{-1}$ | Dilution rate | Experimental set-up: 1 volume change every 8 hours, so 3 per day. The dilution rate and local perturbations thereof (i.e., each integration, for each population) were pulled from a normal distribution: $r_i \sim N(3, 0.02^2)$ |
| $K$ | $[OD]$ | Universal or system-wide maximum density in bioreactor | Co-culture final max OD post antibiotic treatment (od_ph_bioreactor.xlsx), pulled from a normal distribution: $K \sim N(\mu(K), \sigma(K)^2)$. |
| $K_{cd}$ | $[OD]$ | Species-specific maximum density of *C. diff* | Monoculture max OD for *C. diff* (mBhi_GC.xlsx), pulled from a normal distribution: $K_{cd} \sim N(\mu(K_{cd}), \sigma(K_{cd})^2)$. |
| $\alpha_{i,j}$ | $[OD]^{-1}$ | Effect of species $j$ on per capita growth rate of species $i$ | The interaction coefficients were all made relative to the subjected species' growth rate, $r_i$, so that one unit of the exerting species, $j$, corresponds to a fraction, $f$, of the subjected species' growth rate being impacted, and thus $\alpha_{i,j} = f \cdot r_i$. If the direction of interaction (i.e., positive, or negative) was inferable from experimental data, the relative strengths of interaction ($f$) were pulled from uniform distributions following $f \sim U(0.01, 1)$, otherwise: $f \sim N(0, 0.2^2)$ (assuming neutrality, with mean of 0). |
| $x$ | unitless | Multiplier capturing the community-level sensitivity to the antibiotic clindamycin, added between days 8-11, in system between days 8-12. | Relative loss in abundance, estimated from drop in relative abundance during antibiotics treatment. 0 for complete bacteriostatic response, 1 for complete insensitivity. Based on the average relative abundance losses observed, 0.6 was assigned for the multiplier of *E. coli*, and 0 for the multiplier of *F. saccharivorans*. All other species variables had a multiplier value of 1 (no evident community-level sensitivity to the antibiotics). |
| **Initial conditions** | $[OD]$ | Initial density for each of the modelled populations. | Since the OD at t0 of the monoculture growth experiments (mBhi_GC.xlsx) was $\sim 0.01$, all variables were initialised at 0.01 OD, except *C. difficile*. *C. difficile* was initialised with 0.01 OD at day 13 (when it is introduced in the experimental timeline). This introduction density is small enough to warrant an introduction load of an invasive/emergent pathogen, yet big enough to exclude stochastic extinction in the used dilution regime. |

**Supplementary Table 3 B:** Justifications of constraints for interaction parameter inference as used in Ecological Modelling (see Methods 4.2.2.)

| Effect from | Effect to | Direction inferred | Justification | Data source(s) for justification |
| --- | --- | --- | --- | --- |
| *B. caccae* | *E. coli* | positive | With co-cultures achieving mean maximum OD values of >6 (Suppl. fig. x), and *E. coli* achieving >50% of final relative abundance in community co-culture – both when *F. saccharivorans* was its fellow top-abundant member or when this niche was filled by *B. caccae* – approximately >3 community OD units are attributable to *E. coli*, which is much greater than the 1 ODU at which *E. coli* equilibrates in monoculture. Hence, post-antibiotic perturbation, one can deduce that *E. coli* engaged in beneficial (indirect) interactions with other (dominant) Mix-14 members, in particular *B. caccae*. | Mix-14 Community OD data, Mix-14 relative abundance data |
| *E. coli* | *B. caccae* & *F. saccharivorans* | neutral to positive | Vice versa, while *E. coli* is a dominant member of the community, it does not competitively exclude others and dominate the co-culture in its entirety. In fact, GEM results indicate this species may play a 'central donor' role, with the potential to supply metabolites other Mix-14 members, including *B. caccae*, are predicted to be auxotrophic for (Fig. 3a). Dynamically, *F. saccharivorans*' anticorrelative behaviour with *E. coli* around system equilibrium in the absence of/before AB, and *B. caccae*'s sharp rise in relative abundance during/after antibiotics treatment would not be biologically feasible if *E. coli* exerted an antagonistic effect on *B. caccae*. | Mix-14 GEM results, Mix-14 relative abundance data |
| SEC-species & *B. caccae* | *C. difficile* | negative | As shown in individual suppressiveness data (Fig. 1b), SEC-cluster members and *B. caccae* show strong individual suppressive capacities. Moreover, all of these species perform relatively better in post-antibiotic treated and suppressive Mix-14. Hence, they are considered as major players directly or indirectly inhibiting *C. difficile*. In addition, BIOLOG carbon utilisation assay showed close clustering of *C. difficile* with *B. caccae*, implying competition as a potential driver of their interaction. | Individual suppressiveness, *C. difficile* CFU counts at Mix-14 experimental endpoints, relative abundance data, BIOLOG carbon utilisation assay |
| *F. saccharivorans* | *B. caccae* & SEC-species | negative | *F. saccharivorans* and *B. caccae* occupy a near-identical dynamic niche in Mix-14 across experimental conditions, and only occupy this when the other is in low abundance (e.g., *F. saccharivorans* occupies this niche and maintains this position until perturbation by antibiotics or *C. difficile*, after which *F. saccharivorans* declines to near-zero relative abundance and *B. caccae* rises out of low-abundance to occupy the same dynamic niche as a top-abundant species, anti-correlating around system equilibrium with *E. coli*). In addition, in the BIOLOG carbon utilisation data, both of these species cluster closely together, hinting at potential competition between these species). | BIOLOG carbon utilisation assay, Mix-14 relative abundance data |
| *B. caccae* | *F. saccharivorans* | negative | *F. saccharivorans* and *B. caccae* occupy a near-identical dynamic niche in Mix-14 across experimental conditions, and only occupy this when the other is in low abundance (e.g., *F. saccharivorans* occupies this niche and maintains this position until perturbation by antibiotics or *C. difficile*, after which *F. saccharivorans* declines to near-zero relative abundance and *B. caccae* rises out of low-abundance to occupy the same dynamic niche as a top-abundant species, anti-correlating around system equilibrium with *E. coli*). In addition, in the BIOLOG carbon utilisation data, both of these species cluster closely together, hinting at potential competition between these species). | BIOLOG carbon utilisation assay, Mix-14 relative abundance data |
| *B. caccae* | SEC-species | positive | As *B. caccae* rises following antibiotics treatment, so do SEC-species. Since the SEC-species 'follow' the rise of *B. caccae*, the dynamic resembles a commensalist or mutualist (net) interaction between these species groups, and would not be congruent with an antagonistic effect of *B. caccae* on the SEC-species. | Mix-14 relative abundance data |
| SEC-species | *B. caccae* | neutral to positive | As *B. caccae* rises following antibiotics treatment, so do SEC-species. Since the SEC-species 'follow' the rise of *B. caccae*, the dynamic resembles a commensalist or mutualist (net) interaction between these species groups, and would not be congruent with an antagonism between the SEC-species and *B. caccae*. | Mix-14 relative abundance data |
| *C. difficile* | *B. caccae* | neutral to positive | Positive transient effect observed of *C. difficile* on *B. caccae*'s relative abundance (when comparing Mix-14 + AB to Mix-14 + AB + *C. difficile*, and/or when comparing Mix-14 control to Mix-14 + *C. difficile*) | Mix-14 relative abundance data |

## Supplementary Methods for Untargeted Metabolomics

Briefly, the protocol used for analysis of the metabolomics data, as described by Metabolon [88], was as follows:

### Sample Accessioning

Following receipt, samples were inventoried and immediately stored at −80°C. Each sample received was accessioned into the Metabolon LIMS system and was assigned by the LIMS a unique identifier that was associated with the original source identifier only. This identifier was used to track all sample handling, tasks, results, etc. The samples (and all derived aliquots) were tracked by the LIMS system. All portions of any sample were automatically assigned their own unique identifiers by the LIMS when a new task was created; the relationship of these samples was also tracked. All samples were maintained at −80°C until processed.

### Sample Preparation

Samples were prepared using the automated MicroLab STAR® system from Hamilton Company. Several recovery standards were added prior to the first step in the extraction process for QC purposes. To remove protein, dissociate small molecules bound to protein or trapped in the precipitated protein matrix, and to recover chemically diverse metabolites, proteins were precipitated with methanol under vigorous shaking for 2 min (Glen Mills GenoGrinder 2000) followed by centrifugation. The resulting extract was divided into five fractions: two for analysis by two separate reverse phase (RP)/UPLC-MS/MS methods with positive ion mode electrospray ionization (ESI), one for analysis by RP/UPLC-MS/MS with negative ion mode ESI, one for analysis by HILIC/UPLC-MS/MS with negative ion mode ESI, and one sample was reserved for backup. Samples were placed briefly on a TurboVap® (Zymark) to remove the organic solvent. The sample extracts were stored overnight under nitrogen before preparation for analysis.

### QA/QC

Several types of controls were analyzed in concert with the experimental samples: a pooled matrix sample generated by taking a small volume of each experimental sample (or alternatively, use of a pool of well-characterized human plasma) served as a technical replicate throughout the data set; extracted water samples served as process blanks; and a cocktail of QC standards that were carefully chosen not to interfere with the measurement of endogenous compounds were spiked into every analyzed sample, allowed instrument performance monitoring and aided chromatographic alignment. Tables below describe these QC samples and standards. Instrument variability was determined by calculating the median relative standard deviation (RSD) for the standards added to each sample before injection into the mass spectrometer. Overall process variability was determined by calculating the median RSD for all endogenous metabolites (i.e., non-instrument standards) present in 100% of the pooled matrix samples. Experimental samples were randomized across the platform run with QC samples spaced evenly among the injections, as outlined in Suppl. Methods Fig. 1.

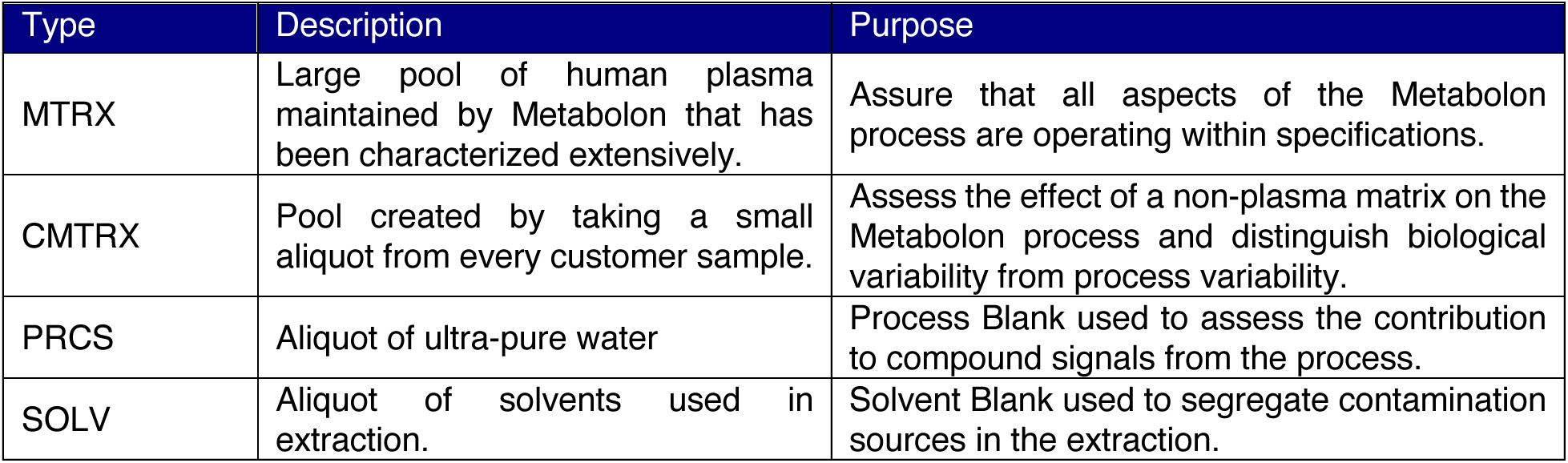

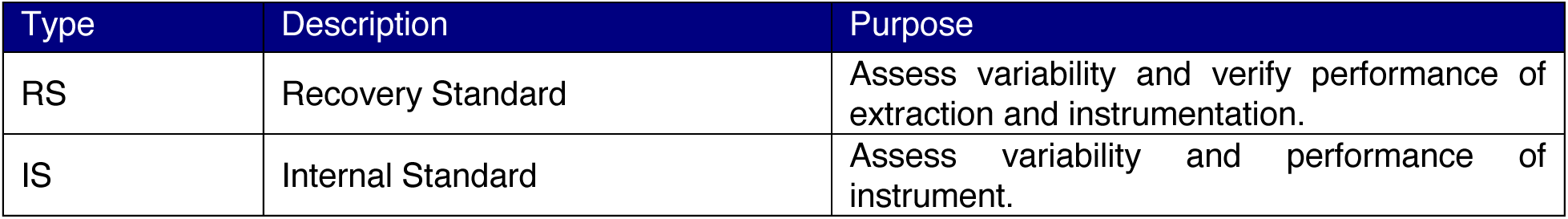

**Supplementary Methods, Fig. 1:**
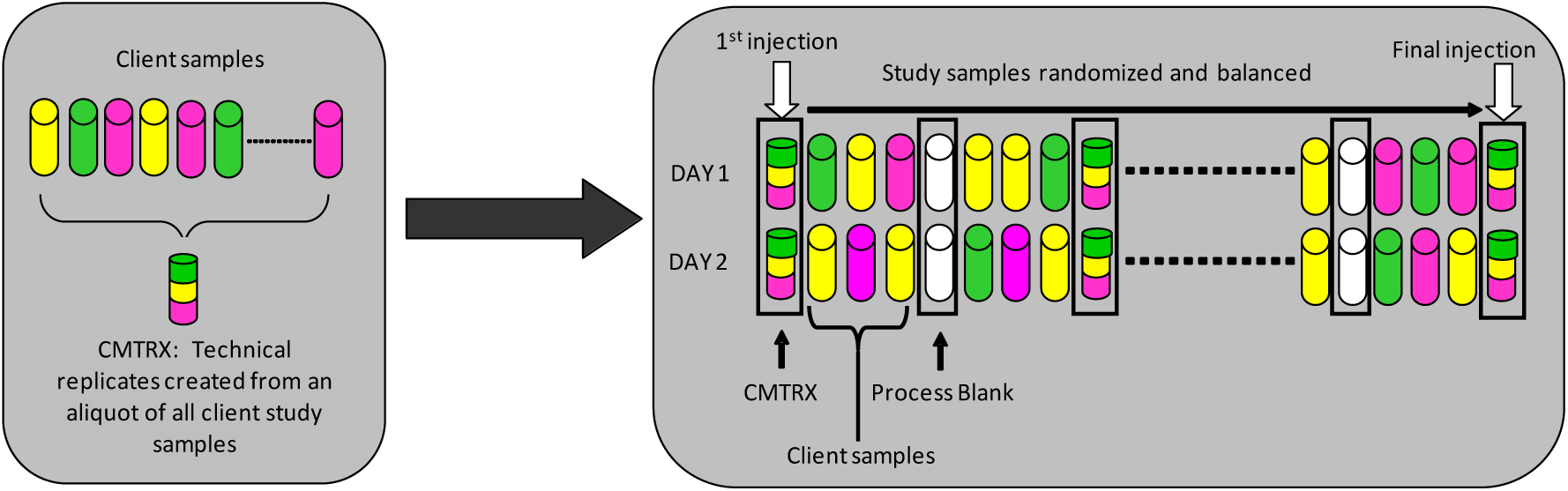
Preparation of client-specific technical replicates. A small aliquot of each client sample (colored cylinders) is pooled to create a CMTRX technical replicate sample (multi-colored cylinder), which is then injected periodically throughout the platform.

### Ultrahigh Performance Liquid Chromatography-Tandem Mass Spectroscopy (UPLC-MS/MS)

All methods utilized a Waters ACQUITY ultra-performance liquid chromatography (UPLC) and a Thermo Scientific Q-Exactive high resolution/accurate mass spectrometer interfaced with a heated electrospray ionization (HESI-II) source and Orbitrap mass analyzer operated at 35,000 mass resolution. The sample extract was dried then reconstituted in solvents compatible to each of the four methods. Each reconstitution solvent contained a series of standards at fixed concentrations to ensure injection and chromatographic consistency. One aliquot was analyzed using acidic positive ion conditions, chromatographically optimized for more hydrophilic compounds. In this method, the extract was gradient eluted from a C18 column (Waters UPLC BEH C18-2.1x100 mm, 1.7 µm) using water and methanol, containing 0.05% perfluoropentanoic acid (PFPA) and 0.1% formic acid (FA). Another aliquot was also analyzed using acidic positive ion conditions, however it was chromatographically optimized for more hydrophobic compounds. In this method, the extract was gradient eluted from the same afore mentioned C18 column using methanol, acetonitrile, water, 0.05% PFPA and 0.01% FA and was operated at an overall higher organic content. Another aliquot was analyzed using basic negative ion optimized conditions using a separate dedicated C18 column. The basic extracts were gradient eluted from the column using methanol and water, however with 6.5mM Ammonium Bicarbonate at pH 8. The fourth aliquot was analyzed via negative ionization following elution from a HILIC column (Waters UPLC BEH Amide 2.1x150 mm, 1.7 µm) using a gradient consisting of water and acetonitrile with 10mM ammonium formate, pH 10.8. The MS analysis alternated between MS and data-dependent MS^n^ scans using dynamic exclusion. The scan range varied slighted between methods but covered 70-1000 m/z. Raw data files are archived and extracted as described below.

### Bioinformatics

The informatics system consisted of four major components, the Laboratory Information Management System (LIMS), the data extraction and peak-identification software, data processing tools for QC and compound identification, and a collection of information interpretation and visualization tools for use by data analysts. The hardware and software foundations for these informatics components were the LAN backbone, and a database server running Oracle 10.2.0.1 Enterprise Edition.

### LIMS

The purpose of the Metabolon LIMS system was to enable fully auditable laboratory automation through a secure, easy to use, and highly specialized system. The scope of the Metabolon LIMS system encompasses sample accessioning, sample preparation and instrumental analysis and reporting and advanced data analysis. All the subsequent software systems are grounded in the LIMS data structures. It has been modified to leverage and interface with the in-house information extraction and data visualization systems, as well as third party instrumentation and data analysis software.

### Data Extraction and Compound Identification

Raw data was extracted, peak-identified and QC processed using Metabolon’s hardware and software. These systems are built on a web-service platform utilizing Microsoft’s .NET technologies, which run on high-performance application servers and fiber-channel storage arrays in clusters to provide active failover and load-balancing. Compounds were identified by comparison to library entries of purified standards or recurrent unknown entities. Metabolon maintains a library based on authenticated standards that contain the retention time/index (RI), mass to charge ratio (*m/z)*, and chromatographic data (including MS/MS spectral data) on all molecules present in the library. Furthermore, biochemical identifications are based on three criteria: retention index within a narrow RI window of the proposed identification, accurate mass match to the library +/- 10 ppm, and the MS/MS forward and reverse scores between the experimental data and authentic standards. The MS/MS scores are based on a comparison of the ions present in the experimental spectrum to the ions present in the library spectrum. While there may be similarities between these molecules based on one of these factors, the use of all three data points can be utilized to distinguish and differentiate biochemicals. More than 3300 commercially available purified standard compounds have been acquired and registered into LIMS for analysis on all platforms for determination of their analytical characteristics. Additional mass spectral entries have been created for structurally unnamed biochemicals, which have been identified by virtue of their recurrent nature (both chromatographic and mass spectral). These compounds have the potential to be identified by future acquisition of a matching purified standard or by classical structural analysis.

### Curation

A variety of curation procedures were carried out to ensure that a high quality data set was made available for statistical analysis and data interpretation. The QC and curation processes were designed to ensure accurate and consistent identification of true chemical entities, and to remove those representing system artifacts, mis-assignments, and background noise. Metabolon data analysts use proprietary visualization and interpretation software to confirm the consistency of peak identification among the various samples. Library matches for each compound were checked for each sample and corrected if necessary.

### Metabolite Quantification and Data Normalization

Peaks were quantified using area-under-the-curve. For studies spanning multiple days, a data normalization step was performed to correct variation resulting from instrument inter-day tuning differences. Essentially, each compound was corrected in run-day blocks by registering the medians to equal one (1.00) and normalizing each data point proportionately (termed the “block correction”; Suppl. Methods Fig. 2). For studies that did not require more than one day of analysis, no normalization is necessary, other than for purposes of data visualization. In certain instances, biochemical data may have been normalized to an additional factor (e.g., cell counts, total protein as determined by Bradford assay, osmolality, etc.) to account for differences in metabolite levels due to differences in the amount of material present in each sample.

**Supplementary Methods, Fig. 2:**
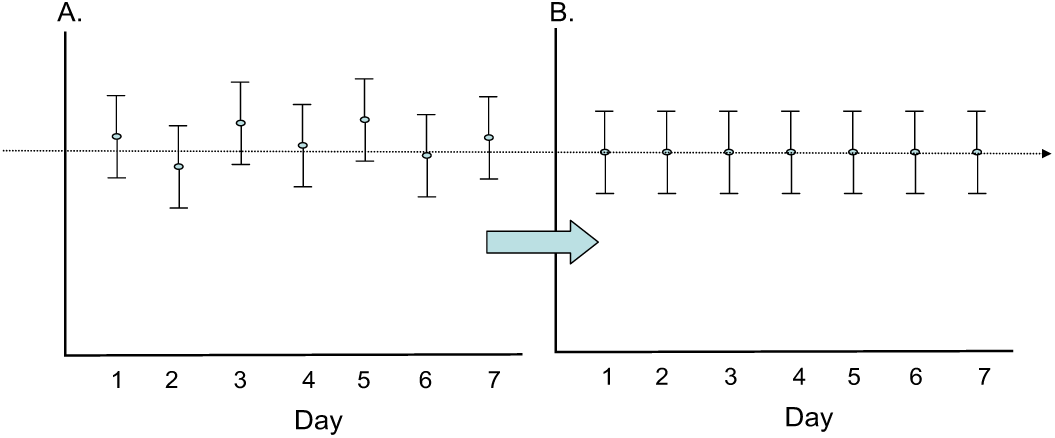
Visualization of data normalization steps for a multiday platform run.

